# Multi-level transcriptional regulation of embryonic sex determination and dosage compensation by the X-signal element *sex-1*

**DOI:** 10.1101/2024.11.23.624987

**Authors:** Eshna Jash, Zoey M. Tan, Audry I. Rakozy, Anati Alyaa Azhar, Hector Mendoza, Györgyi Csankovszki

## Abstract

The *C. elegans* nuclear hormone receptor *sex-1* is known to be an embryonic X-signal element that represses *xol-1*, the sex-switch gene that is the master regulator of sex determination and dosage compensation. Several prior studies on *sex-1* function have suggested that *sex-1* may have additional downstream roles beyond the regulation of *xol-1* expression. In this study we characterize some of these additional roles of *sex-1* in regulating the dual processes of sex determination and dosage compensation during embryogenesis. Our study reveals that *sex-1* acts on many of the downstream targets of *xol-1* in a *xol-1*-independent manner. Further analysis of these shared but independently regulated downstream targets uncovered that *sex-1* mediates the expression of hermaphrodite- and male-biased genes during embryogenesis. We validated *sex-1* binding on one of these downstream targets, the male-developmental gene *her-1.* Our data suggests a model where *sex-1* exhibits multi-level direct transcriptional regulation on several targets, including *xol-1* and genes downstream of *xol-1,* to reinforce the appropriate expression of sex-biased transcripts in XX embryos. Furthermore, we found that *xol-1 sex-1* double mutants show defects in dosage compensation. Our study provides evidence that misregulation of *dpy-21*, one of the components of the dosage compensation complex, and the subsequent misregulation of H4K20me1 enrichment on the X chromosomes, may contribute to this defect.

## Introduction

In the nematode *C. elegans*, sex is determined by the number of X chromosomes relative to autosomes inherited by the fertilized embryo from parental gametes. Hermaphrodites have two X chromosomes, and males only have one X chromosome to two sets of autosomes. This differential inheritance of sex chromosomes in the two sexes kick-starts a molecular cascade in the developing embryo that results in the formation of the appropriate sexual morphology, gonadogenesis and dosage compensation. The molecular mechanisms of sex determination in *C. elegans* depends on the ratio of X chromosomes to autosomes (X:A ratio) [1], [2]. Hermaphrodites have a higher X:A ratio of 1 (XX:AA), and males have a lower X:A ratio of 0.5 (X:AA). XOL-1 is the “sex-switch” gene that is sensitive to this difference in X:A ratio, and acts as the master regulator of sex determination and dosage compensation [3], [4] (Fig. 1A).

**Fig. 1:**
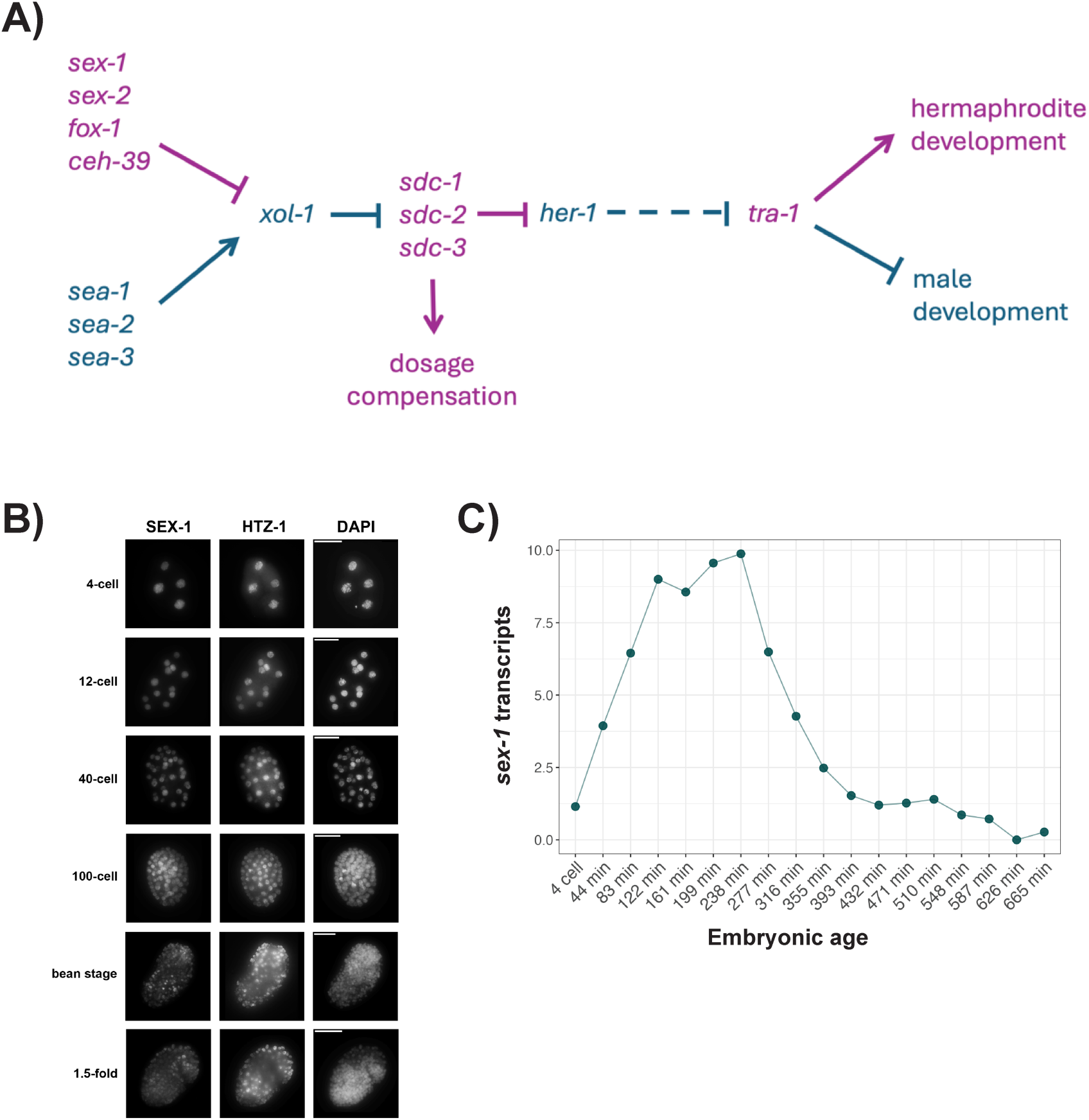
Patterns of *sex-1* expression and localization. (A) Schematic depicting the canonical pathway regulating sex determination and dosage compensation in *C. elegans* embryogenesis. Genes that promote hermaphrodite-specific development are indicated in purple and genes that promote male-specific development are indicated in blue. (B) Immunofluorescence staining with anti-FLAG antibodies to determine localization patterns of SEX-1::FLAG in embryos. Antibodies against the histone H2A variant HTZ-1 were used as staining control. (C) Scatter plot depicting pattern of *sex-1* transcript expression in embryos using time-resolved embryogenesis transcriptomic dataset [34].

XOL-1 is structurally homologous to GHMP kinases and belongs to a unique subclass within the GHMP kinase family [5]. XOL-1 acts through the SDC proteins to regulate the independent pathways of sex determination and dosage compensation [6]. Activation of the SDC proteins SDC-1, SDC-2 and SDC-3 results in the transcriptional repression of a crucial male-developmental gene *her-1* [7], [8], [9]. *her-1* is a potent activator of the male sex determination pathway [7]. Expression of *her-1* in embryos is both necessary and sufficient to initiate male-specific somatic differentiation [7], [8]. Somatic cells in *C. elegans* are highly sexually dimorphic, with an estimated ∼30% of somatic cells in post-embryonic cells having sex-specific differences in morphology and function [10].

The SDC proteins also separately regulate the process of dosage compensation [11], [12], [13], [14], [15]. Similar to X chromosomes in other organisms, the *C. elegans* X chromosomes are gene-rich. Therefore, the difference in gene dosage needs to be balanced between the two sexes. This is accomplished through the process of dosage compensation, carried out by the dosage compensation complex (DCC) in *C. elegans* [16], [17], [18], [19]. The DCC is a 10-subunit protein complex consisting of a pentameric condensin I^DC^ complex and five additional proteins which includes the three SDC proteins [17], [19], [20], [21]. The DCC is recruited to the X chromosomes by these SDC proteins [11], [16], [18]. The DCC compacts the X chromosomes through the chromatin looping activity of condensin I^DC^, a *C. elegans*-specific condensin complex, and through the additional recruitment of several pathways that reinforce this X repression [22]. Additional pathways that have been characterized include the deposition of repressive epigenetic marks such as H4K20me1 by the DCC subunit DPY-21 [23], [24], [25], and H3K9me3-mediated tethering of the X chromosomes to the nuclear lamina [26].

Beyond the canonical roles of *xol-1* in promoting the male sex determination pathway through repression of the activity of the SDC proteins, XOL-1 also has important roles in regulating events during early embryogenesis in XX hermaphrodites [27]. In a recent study we showed that low-level expression of XOL-1 during early embryonic development in hermaphrodites is important for regulating the speed of embryonic development, as well as the timing of dosage compensation initiation on the X chromosomes [27]. *xol-1* mutant embryos appear to have accelerated embryonic development in terms of cell division and activation of embryonic stage-specific transcriptional programs [27]. Furthermore, the DCC loads onto the X chromosomes at earlier stages of embryonic development upon loss of *xol-1* [27].

XOL-1 “senses” the X:A ratio in an embryo through the action of upstream regulators that are transcribed from the X chromosomes and the autosomes. The regulators transcribed from the X chromosomes are known as X-signal elements (XSEs) and those transcribed from the autosomes are known as autosomal-signal elements (ASEs) [1], [2]. These regulators compete to repress and activate XOL-1, respectively. XSEs and ASEs function in a dose-dependent, cumulative manner so that the difference in the dosage of X chromosomes results in either the activation of XOL-1 in the case of XO males, or its repression in the case of XX hermaphrodites. Several of the elements have been characterized, such as the XSEs *sex-1*, *sex-2*, *fox-1* and *ceh-39*, and ASEs such as *sea-1*, *sea-2* and *sea-3* [1], [28], [29], [30], [31], [32]. These elements have several different mechanisms of action. XSEs such as *sex-1* and *ceh-39* bind directly to specific sites on the *xol-1* promoter region to transcriptionally repress the *xol-1* gene [1]. *fox-1*, on the other hand, is a splicing regulator that alters the splicing of the *xol-1* pre-mRNA, resulting in a non-functional XOL-1 protein [30]. In a mechanism similar to *sex-1* and *ceh-39*, the ASE *sea-1* directly binds to sites on the *xol-1* promoter to activate its transcription [1]. The mechanism of *xol-1* regulation by the ASE *sea-2* is also through transcriptional activation of the *xol-1* gene. *sea-2* was shown to activate *xol-1* transcription using an *in vivo* reporter containing *xol-1* promoter fragments [1]. *sea-2* has also been shown to be capable of binding to RNA fragments *in vitro*, suggesting that it may post-transcriptionally regulate *xol-1* activation as well [32].

Though the XSEs act in an additive manner, the contribution of each XSE towards *xol-1* repression is not equivalent. Mutational and epistasis analysis of XSEs and ASEs in combination has revealed a hierarchy in XSE strength [2]. Among the XSEs, *sex-1* has the biggest effect on the repression of *xol-1* [2]. SEX-1 is a transcription factor that belongs to the nuclear hormone receptor (NHR) superfamily [28]. SEX-1 is a transcriptional repressor of XOL-1 during the early stages of embryogenesis, as evidenced by *in vivo* assays [28]. Additionally, *in vitro* assays have demonstrated that SEX-1 binds to NHR consensus sites in the promoter of the *xol-1* gene [1]. SEX-1 is homologous to NHRs in other organisms such as the *Drosophila* ecdysone-sensitive NHR E78A and the human NHR Rev-erb [28]. The DNA-binding domain of SEX-1 is highly similar to other NHRs, which allows it to bind to the NHR consensus sequences [1], [28]. However, other domains of the SEX-1 protein have diverged from its closely related proteins. SEX-1 has no known ligands and a highly divergent ligand-binding domain (LBD) [28].

Though SEX-1 has well-characterized roles in relaying X dosage for sex determination during early embryogenesis, it has long been known that SEX-1 may have additional roles beyond transcriptional regulation of *xol-1*. Several prior studies have provided evidence for a *xol-1*-independent role in both sex determination and dosage compensation [2], [33]. *sex-1* null mutants are very unhealthy, with severe embryonic and larval lethality. These phenotypes are hypothesized to be caused by the ectopic expression of high levels of *xol-1* in hermaphrodite embryos. However, *xol-1 sex-1* double null mutants are not able to completely rescue *sex-1*-mediated lethality, indicating that *sex-1* has additional roles beyond *xol-1* regulation [2]. Another line of evidence for a role of *sex-1* downstream of *xol-1* comes from male rescue experiments. *xol-1* deletion causes male-specific lethality due to the activation of the dosage compensation machinery on the single male X. *xol-1 sex-1* mutants show rescue of male lethality phenotype, suggesting that *sex-1* acts independent of *xol-1* regulation to promote dosage compensation [2], [33]. Disrupting this function leads to a male rescue phenotype by destabilizing the activation of dosage compensation on the male X chromosome. Furthermore, some of these rescued XO animals develop as phenotypic hermaphrodites, suggesting that loss of *sex-1* disrupts the pathways of sex determination as well [2], [33].

In this paper we characterize the downstream roles of *sex-1* that are independent of its regulation of *xol-1*. Our data suggests that *sex-1* antagonizes the *xol-1* pathway by directly or indirectly acting on targets downstream of *xol-1* that function in the processes of sex determination, dosage compensation and timing of embryonic development. We provide evidence that *sex-1* regulates the male sex determination pathway through direct transcriptional regulation of the male developmental gene *her-1*. Additionally, we provide evidence that *sex-1* mediates dosage compensation through regulation of the repressive H4K20me1 mark deposited by the DCC subunit DPY-21. Our data also suggests that the disruption of dosage compensation mediated by H4K20me1 is likely not the cause of decreased viability in *xol-1 sex-1* double mutants.

## Results

### Patterns of *sex-1* expression and localization

To understand the function of SEX-1, we first sought to examine its expression patterns within the embryo. To examine the expression of SEX-1 in embryonic tissues, we used CRISPR-Cas9 based gene editing to generate an endogenously tagged SEX-1::FLAG strain. We performed embryonic viability and larval viability quantification on our SEX-1::FLAG strain to confirm that SEX-1 function is not compromised (Fig. S1A-B). While loss of function mutations in *sex-1* lead to reduced viability [2], [28], the viability of *sex-1::FLAG* embryos and larvae were comparable to wild type (Fig. S1A-B)). We then used immunofluorescence staining of embryos with anti-FLAG antibodies to determine where SEX-1 is being expressed in the embryo (Fig. 1B). We observed ubiquitous expression of SEX-1 in embryonic nuclei during the early stages of embryogenesis (Fig. 1B). SEX-1 signal is first visible at the 4-cell stage, where it begins being enriched on some nuclei (Fig. 1B). Clear nuclear enrichment on all nuclei is evident from the 8-cell stage to the 100-cell stage (Fig. 1B). SEX-1 signal gradually tapers off after the bean stage, but remains visible in nuclei throughout embryogenesis (Fig. 1B). These results are largely consistent with what was reported previously for SEX-1 expression using anti-SEX-1 antibodies [28]. SEX-1 protein expression closely follows the patterns of expression of the *sex-1* transcripts during embryonic development. We used a time-resolved transcriptome of embryogenesis to look at stage-specific expression of *sex-1* [34] (Fig. 1C). *sex-1* transcripts are detectable at a low-level in 4-cell embryos, and rapidly accumulate during early embryogenesis (Fig. 1C). Peak transcript levels are detectable at 235 min post fertilization, which corresponds to mid-gastrulation stage (Fig. 1C). After this stage, *sex-1* expression tapers off, but transcripts continue being detected until ∼500min post fertilization, corresponding to late 2-fold embryo stage (Fig. 1C).

### *xol-1 sex-1* partially rescues *sex-1* phenotypes

Several prior studies have hinted at additional roles of *sex-1* in the processes of dosage compensation and sex determination. However, these studies used several mutants where the precise molecular nature of the mutations were uncharacterized and/or used RNAi treatment to disrupt the function of the *xol-1* and *sex-1* genes. To probe the additional downstream functions of *sex-1*, we first obtained well-characterized null mutants for both *xol-1* and *sex-1*. In this study we use the *xol-1(ne4472)* allele first characterized in Tang et al. (2018) [35]. This allele contains frameshift in-del mutations in the first exon of *xol-1*. We derived the *sex-1(gk829)* null allele from a strain generated by the international *C. elegans* Gene Knockout Consortium [36]. This allele contains a large deletion starting at the transcription state site (TSS), and spanning the first and second exons of the *sex-1* gene. For the remainder of this manuscript, these characterized and predicted null alleles will be referred to as *xol-1* and *sex-1* mutants.

To explore the possible *xol-1*-independent roles of *sex-1*, we created a *xol-1 sex-1* double null mutant. A prior study by our lab performed an XO male rescue assay using this *xol-1 sex-1* strain, which confirmed that the introduction of the *sex-1* null allele in a *xol-1* null background is able to partially rescue the male lethality phenotype of *xol-1* mutants [33]. This data suggests that *sex-1* has a role downstream of *xol-1* promoting dosage compensation, confirming the results reported in the Gladden et al. (2007) study [2]. We sought to further examine if our *xol-1 sex-1* strain is able to rescue the various phenotypes observed in the *sex-1* mutant. First, we quantified embryonic viability in WT, *xol-1*, *sex-1* and *xol-1 sex-1* mutants (Fig. 2B). In this assay we quantified the proportion of embryos laid by young adults that are able to survive embryogenesis and hatch into L1 larvae. WT and *xol-1* mutants appear very similar, with no statistically significant differences in embryonic viability (Fig. 2B). *sex-1* mutants appear very unhealthy, with high levels of embryonic lethality (Fig. 2B). These XX hermaphrodite worms inappropriately express *xol-1* during embryonic development, resulting in widespread disruptions of the sex determination and dosage compensation pathways, and ultimately leading to lethality (Fig. 2A). The *xol-1* mutation is able to significantly rescue some, but not all, of the embryonic lethality caused by *sex-1* deletion in *xol-1 sex-1* double mutants (Fig. 2B). Only ∼70% of *sex-1* mutant embryos survive embryogenesis, but the additional loss of *xol-1* in *xol-1 sex-1* double mutant rescues the embryo survival rate to ∼83%. However, embryonic viability in *xol-1 sex-1* strain remains statistically significantly different from both WT and *xol-1* mutant (Fig. 2B, Table S1).

**Fig. 2:**
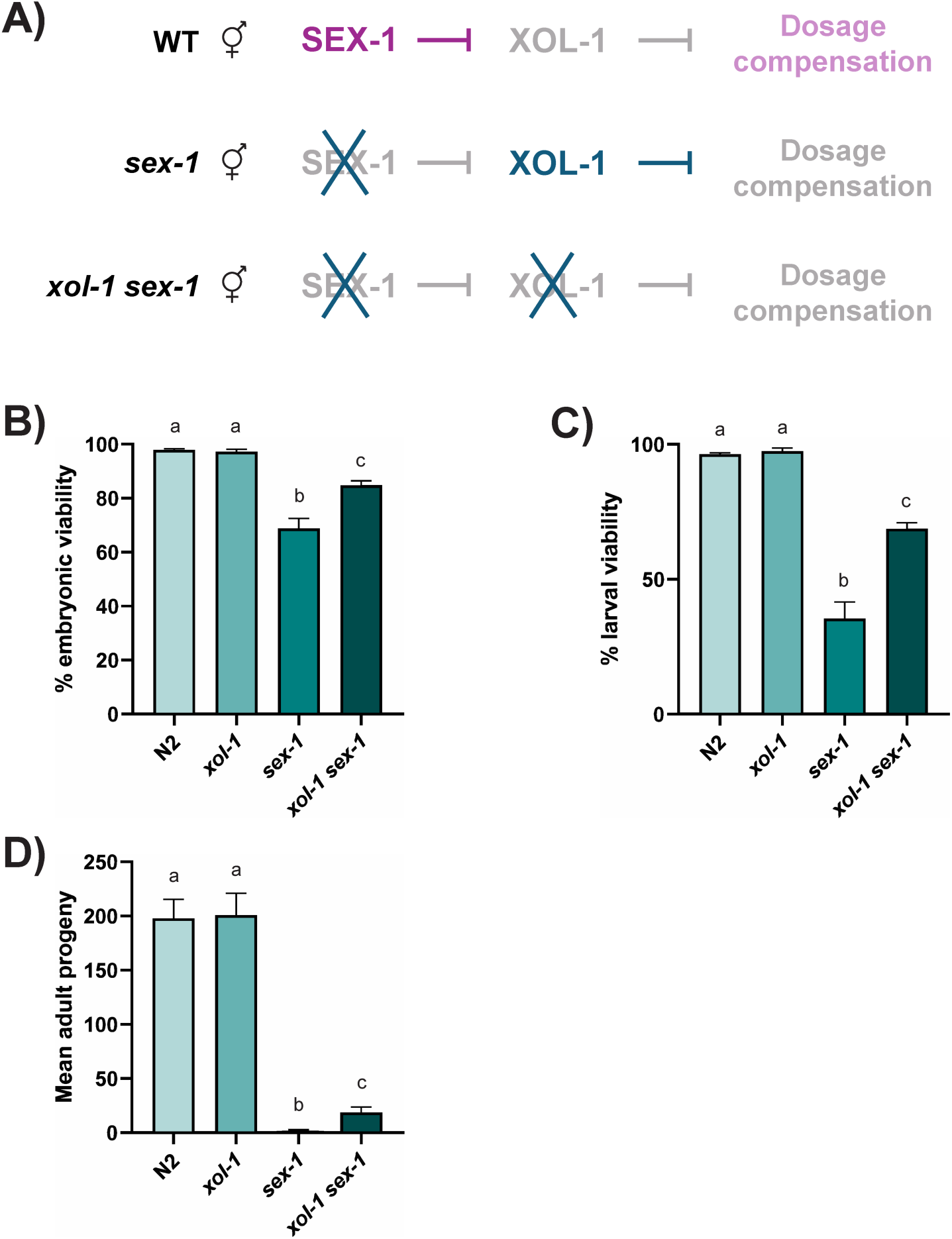
*xol-1 sex-1* partially rescues *sex-1* phenotypes. (A) Schematic depicting the expected activation or inhibition of dosage compensation in WT, *xol-1* and *xol-1 sex-1* strains. Genes that promote hermaphrodite-specific development are indicated in purple and genes that promote male-specific development are indicated in blue. (B) Embryonic viability scored in WT, *xol-1*, *sex-1* and *xol-1 sex-1* strains. P-values obtained from unpaired Welch’s t-test with unequal variance. Error bars indicate SEM. At least 1100 embryos were counted in total from a minimum of 12 biological replicates per genotype. (C) Larval viability scored in WT, *xol-1*, *sex-1* and *xol-1 sex-1* strains. P-values obtained from unpaired Welch’s t-test with unequal variance. Error bars indicate SEM. At least 550 larvae were counted in total from a minimum of 5 biological replicates per genotype. (C) Total viable adult progeny 72h post egg laying scored in WT, *xol-1*, *sex-1* and *xol-1 sex-1* strains. P-values obtained from unpaired Welch’s t-test with unequal variance. Error bars indicate SEM. Progeny from 10 biological replicates were quantified per genotype. Letters above the bars indicate statistical significance (same letters indicate that datasets do not have a statistically significant difference, different letters indicate that datasets have a statistically significant difference). P-values are reported in Table S1.

A similar pattern is seen in the larval viability assay. In this assay we measured the proportion of L1 larvae that develop into young adults in 72 hours. The *xol-1* mutation partially rescues the *sex-1* null phenotype but larval viability in *xol-1 sex-1* double mutants remains significantly below WT and *xol-1* mutants (Fig. 2C). ∼28% of *sex-1* mutant L1 larvae survive larval development to develop into young adults (Fig. 2C). Larval viability in *xol-1 sex-1* mutants is higher at ∼69% but it remains statistically significantly below *xol-1* mutants at ∼97% (Fig. 2C, Table S1). Finally, we measured the total live adult progeny from each of these strains (Fig. 2D, Table S1). *sex-1* mutants have extremely low fecundity with an average of 1.8 live progeny per worm (Fig. 2D). Consistent with prior studies, we also observed that *sex-1* mutant hermaphrodites that do survive embryonic and larval development frequently have a masculinized or intersex phenotype [28]. Similar to the embryonic and larval viability assays, *xol-1* is able to rescue this phenotype, but only partially. *xol-1 sex-1* worms lay on average 18.6 progeny per worm (Fig. 2D). This is far lower than the number of progeny laid by WT and *xol-1* worms that are at ∼200 progeny per worm. Taken together, the data from these assays confirm that *sex-1* has functions beyond that of an X-signal element that represses the transcription of the *xol-1* gene. The loss of the *xol-1*-independent functions of *sex-1* results in significant embryonic and larval lethality, as well as very low fecundity in *xol-1 sex-1* double mutants.

### *sex-1* and *xol-1* independently regulate a shared downstream pathway

Since SEX-1 is a transcription factor, we hypothesized that it is likely able to bind to promoter sequences of genes other than *xol-1* to regulate their transcription as well. We performed mRNA-seq on synchronized WT, *xol-1* and *xol-1 sex-1* mutant early embryos to explore the changes in gene expression caused by loss of *sex-1*. Early embryo samples collected for sequencing represent embryos from 2-cell to bean stage. The WT and *xol-1* datasets were previously analyzed in [27]. Differentially expressed (DE) genes in the *xol-1*/ WT dataset represent the transcriptional changes caused by the loss of *xol-1* function, and DE genes in the *xol-1 sex-1*/ *xol-1* dataset represent changes caused by the loss of *xol-1*-independent *sex-1* function. A volcano plot shows that there are many genes that are differentially expressed present in this *xol-1 sex-1*/ *xol-1* dataset (Fig. 3A), indicating that many genes are regulated (directly or indirectly) by *sex-1*. Furthermore, several lines of evidence from all three of these transcriptional datasets suggested to us that *sex-1* may independently regulate *xol-1* targets. The venn diagram in Fig. 3B shows that a large number of DE genes are shared between *xol-1*/ WT and *xol-1 sex-1*/ *xol-1*. 3639 genes (41% of genes) are differentially expressed in both *xol-1 sex-1*/ *xol-1* and *xol-1*/ WT. To explore the direction and magnitude of differential expression in these genes, we looked at the Log2 fold change distribution of the entire dataset. In Fig. 3C, we examined the subset of 3639 DE genes that are shared between the two datasets. The genes that are differentially expressed (|Log2 Fold Change| > 0.5, adjusted p-value < 0.05) in both datasets are highlighted in dark blue. These genes appear to be strongly anti-correlated in the two datasets, with a Pearson correlation coefficient of -0.91, indicating a strong and statistically significant negative correlation. The genes highlighted in salmon are not statistically significantly different in at least one of the datasets. This analysis demonstrates that most genes that are upregulated in *xol-1*/ WT are downregulated in *xol-1 sex-1/ xol-1*, and vice versa. A very small number of genes (109 genes, 3%) appear to be directly correlated. We removed these genes and ran linear modeling on our dataset to examine the relative magnitude of change in the shared and anti-correlated transcripts. Linear modeling on these shared transcripts confirms this anti-correlation (R = -0.95) with a very tight 95% confidence interval (Fig. 3D). There appears to be a similar degree of change in expression for genes regulated by *xol-1* and by the *xol-1*-independent function of *sex-1*, with slightly greater change in the *xol-1*/ WT dataset. Taken together, this data strongly suggests a model where *sex-1* regulates transcriptional targets downstream of *xol-1* and antagonizes *xol-1* function.

**Fig. 3:**
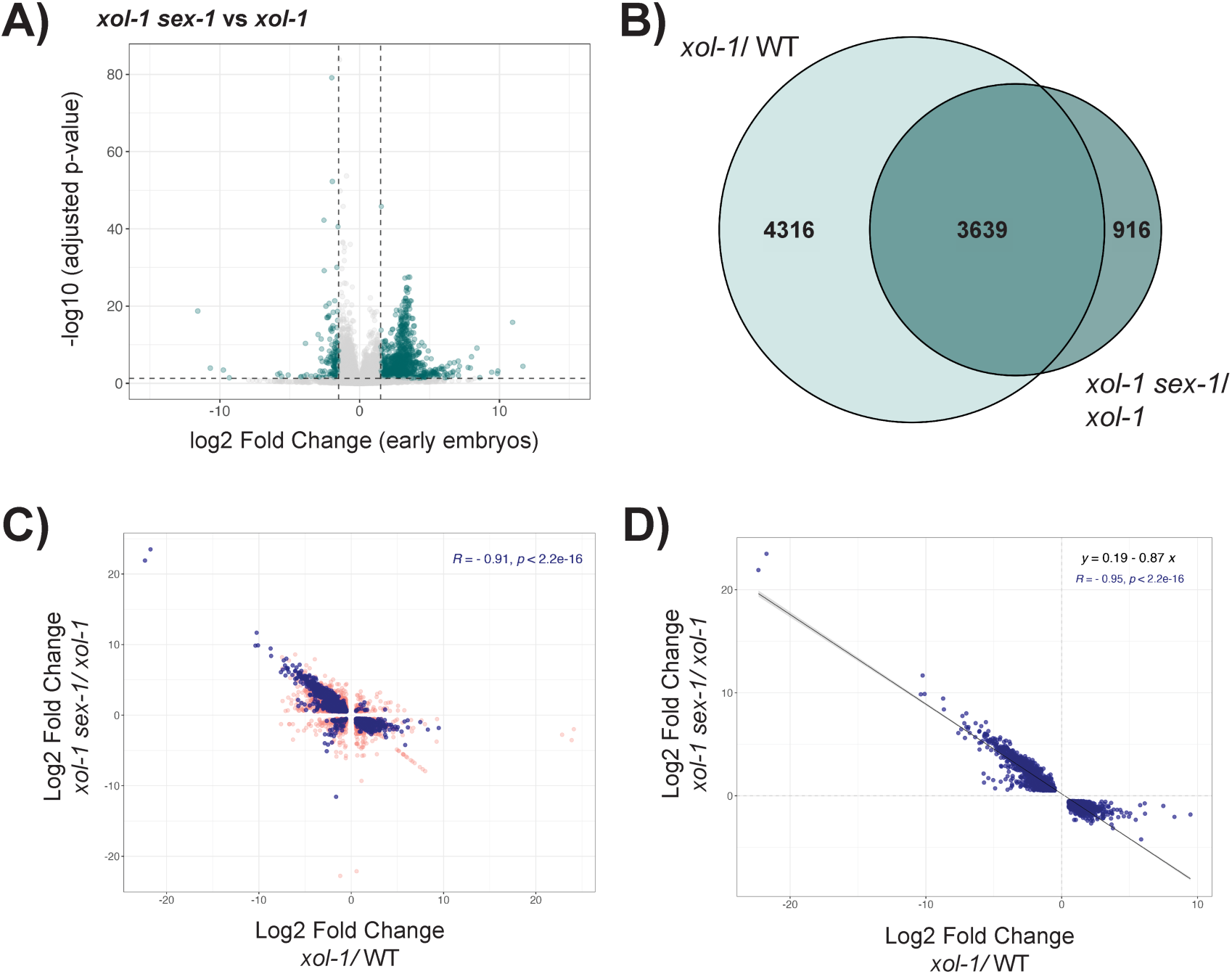
*sex-1* and *xol-1* independently regulate a shared downstream pathway. (A) Volcano plot representing differentially expressed (DE) genes in *xol-1 sex-1*/ *xol-1* early embryo dataset. Dotted lines represent thresholds used to define DE genes: absolute log2 fold change > 1.5 and adjusted p-value < 0.05. (B) Venn diagram representing shared differentially expressed genes (|log2 fold change| > 0.5, adjusted p-value < 0.05) between *xol-1*/ WT and *xol-1 sex-1*/ *xol-1*. (C-D) Scatter plots with log2 fold change in *xol-1 sex-1*/ *xol-1* (y-axis) and *xol-1*/ WT (x-axis) with corresponding Pearson correlation coefficient (R) (C) Blue points represent genes differentially expressed in both datasets, salmon points represent genes that are not statistically significantly different in at least one dataset. (D) Blue points represent genes differentially expressed in both datasets and anti-correlated in their expression pattern between datasets.

### The *sex-1* mutation reverses gene expression changes caused by loss of *xol-1*

In a previous study we showed that loss of *xol-1* in hermaphrodite early embryos results in an acceleration of embryonic development that results in broad changes in the embryonic transcriptional program [27]. Loss of *xol-1* in hermaphrodite embryos resulted in the upregulation of genes expressed in late embryos, and the downregulation of genes expressed in the early embryos. To explore the role of *sex-1* [27], we looked at gene expression changes between genes expressed in early-stage embryos and those expressed in late-stage embryos (Fig. 4A-B). Genes expressed in early embryos appeared to be upregulated in *xol-1 sex-1* mutants compared to *xol-1* mutants, and unchanged in *xol-1 sex-1* mutants when compared to WT embryos (Fig. 4A). Conversely, genes expressed in late-stage embryos are downregulated in *xol-1 sex-1* mutants compared to *xol-1* mutants, and unchanged in *xol-1 sex-1* mutants vs WT (Fig. 4B). As measured by their transcriptional status, *xol-1 sex-1* embryos do not appear to be developmentally accelerated compared to WT. Since *xol-1 sex-1* double mutant reverses the transcriptional changes associated with the embryonic acceleration phenotype seen in *xol-1* embryos, this data implies that *sex-1* is independently acting on *xol-1* targets that regulate developmental timing. *met-2* is overexpressed in *xol-1* mutants and it is partially responsible for the developmental acceleration observed in *xol-1* null embryos [27]. We examined the transcriptional regulation of *met-2* in *xol-1 sex-1* mutants. However, we did not observe any statistically significant change in *met-2* transcript levels between *xol-1 sex-1* and *xol-1* strains (Fig. S1C). This suggests that the *sex-1* mutation reverses some, but not all changes caused by a mutation in *xol-1,* and that the *sex-1*-mediated reversal in gene expression may be due to its regulation of downstream targets other than *met-2*.

**Fig. 4:**
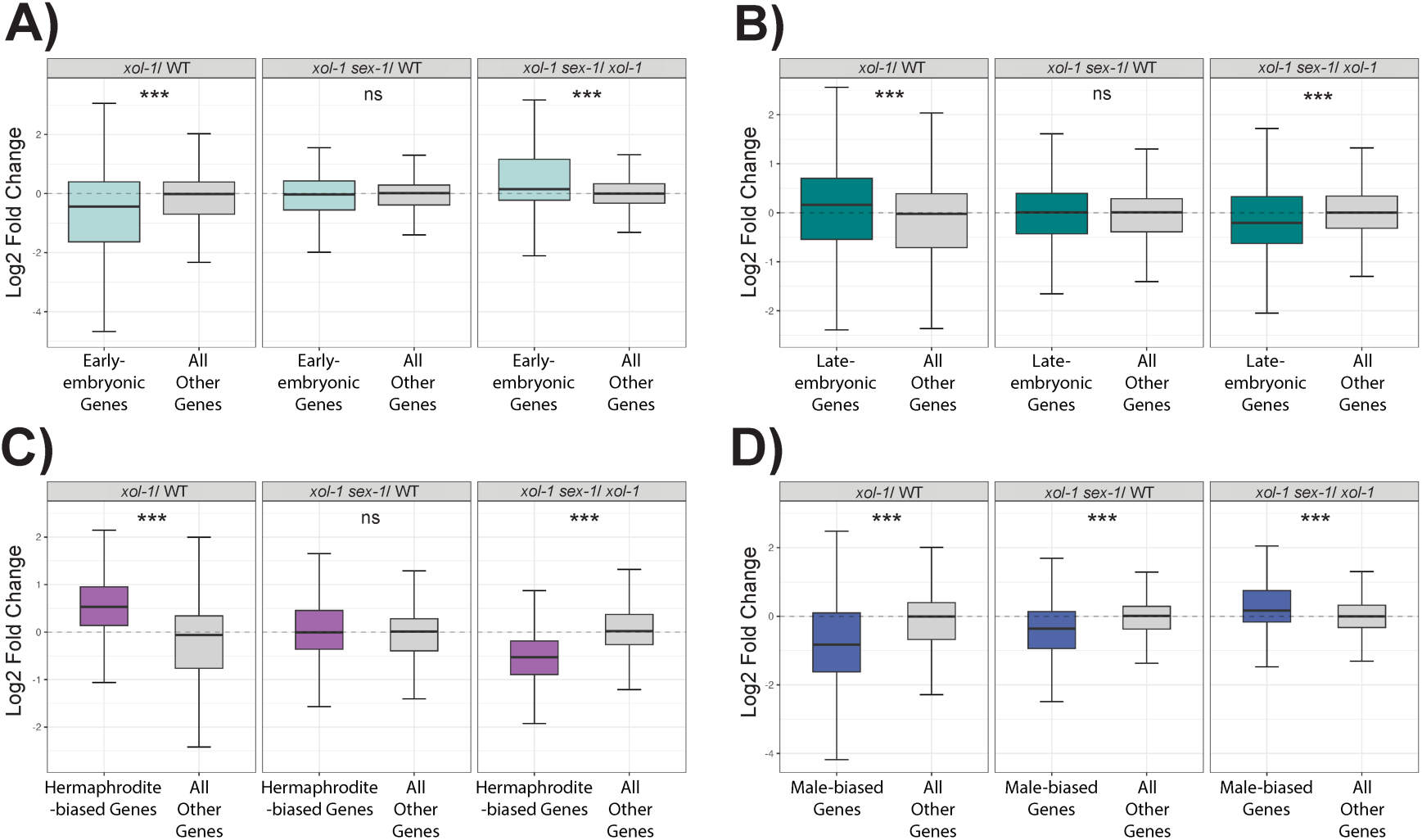
The loss of *sex-1* reverses gene expression changes caused by loss of *xol-1*. (A-D) Boxplots depicting the distribution of log2 fold change in gene expression in *xol-1*/WT, *xol-1 sex-1*/ WT and *xol-1 sex-1*/ *xol-1*. (A) Distribution of log2 fold change in genes known to be expressed in early embryogenesis, obtained from Spencer et al. (2011) [37]. Mood’s median test was used to calculate statistical significance. (B) Genes known to be expressed in late embryogenesis, obtained from Spencer et al. (2011) [37]. Mood’s median test was used to calculate statistical significance. (C) Hermaphrodite-biased genes with correction for altered embryonic development in *xol-1* mutants, obtained from Jash et al. (2024) [27]. Mood’s median test was used to calculate statistical significance. (D) Male-biased genes, with correction for altered embryonic development in *xol-1* mutants, obtained from Jash et al. (2024) [27]. Mood’s median test was used to calculate statistical significance. Asterisks indicate the level of statistical significance (* p < 0.05, ** p < 0.01, *** p < 0.005, ns not significant).

Another consequence of the loss of *xol-1* in hermaphrodites is a disruption in the transcriptional pathways of sex determination during early embryogenesis [27]. This can be measured through the upregulation of hermaphrodite-biased genes and downregulation of male-biased genes in *xol-1* mutant hermaphrodite early embryos [27]. Since we know that *sex-1* also has likely roles in regulating the sex determination pathways downstream of XOL-1 [2], [33], we looked at the gene expression changes in sex-biased transcripts in our *xol-1 sex-1* strain. To do this, we used the *him-8* early embryo dataset we generated in the study described above to define male-biased and hermaphrodite-biased genes [27]. The *him-8* mutation results in ∼40% of the progeny being male, resulting in a male-enriched early embryonic dataset. Genes that were upregulated in this dataset compared to WT were defined as male-biased genes and those that were downregulated were defined as hermaphrodite-biased genes. Furthermore, these gene sets were filtered to account for the difference in embryonic development between *xol-1* mutants and WT, as previously described [27]. In the *xol-1 sex-1* vs *xol-1* dataset, we found the hermaphrodite-biased genes to be significantly downregulated (Fig. 4C). There was no significant difference in the median gene expression of hermaphrodite-biased genes between *xol-1 sex-1* mutants and WT (Fig. 4C). This suggests that in *xol-1 sex-1* mutants the *sex-1* mutation largely reverses the transcriptional changes caused by the *xol-1* mutation to resemble the transcriptional program in WT. We performed a similar analysis to examine the transcriptional status of male-biased genes in these strains. Male-biased transcripts in *xol-1 sex-1* mutants were slightly, but statistically significantly, upregulated compared to *xol-1* mutants. However, male-biased genes still appeared to be downregulated compared to WT (Fig. 4D). This suggests that the *sex-1* mutation in *xol-1 sex-1* double mutants is able to partially rescue the transcriptional changes in male-biased genes, but that these transcripts remain altered due to the loss of *xol-1*.

We confirmed regulation of sex-biased genes by *sex-1* with gene set enrichment analysis (GSEA), a statistical tool to measure the enrichment or depletion of an *a priori* defined set of genes. The gene set enrichment analysis showed that hermaphrodite-biased genes were depleted in *xol-1 sex-1* double mutants compared to *xol-1* mutants, with a normalized enrichment score (NES) of -3.01 (Fig. S2A). Conversely for male-biased genes, the analysis showed an enrichment in *xol-1 sex-1* double mutants compared to *xol-1* mutants, with an NES of 1.64 (Fig. S2A). Despite this statistically significant enrichment, male-biased genes remained depleted when compared to WT, with an NES of -2.17 (Fig. S2A). These data suggest that *sex-1* acts strongly on the hermaphrodite sex determination pathway, and to a lesser extent on the male sex determination pathway in early embryos, in a *xol-1*-independent manner. This data also further strengthens the hypothesis that *sex-1* independently acts on transcriptional targets downstream of *xol-1*.

### *her-1* is a direct transcriptional target of *sex-1*

We sought to confirm this model of multi-level *xol-1* and *sex-1* antagonistic regulation by validating a downstream target of *xol-1* that is also independently regulated by *sex-1*. A candidate for this was reported in a study by Gladden et al. (2007) that showed that the loss of *xol-1*-independent *sex-1* function results in the transcriptional upregulation of *her-1*, a male-developmental gene that is downstream of *xol-1* [2]. *her-1* is also upregulated in our *xol-1 sex-1*/ *xol-1* mRNA-seq dataset (Fig. 5A), confirming the results from Gladden et al. (2007). Additionally, we validated this finding by RT-qPCR measurement of the specific splice isoform of *her-1* that is known to be important for male-specific embryonic development, *her-1a* (Fig. 5B). We also found misregulation of a hermaphrodite-developmental gene, *tra- 1*, in *xol-1 sex-1*/ *xol-1*. *tra-1* is a terminal regulator of sex determination in *C. elegans*, with important roles in repressing male-biased targets and activating the transcription of hermaphrodite-biased targets [38]. *tra-1* is downregulated in *xol-1 sex-1*/ *xol-1* as measured by mRNA-seq and RT-qPCR (Fig. 5A-B), suggesting that *sex-1* promotes the expression of *tra- 1* in a *xol-1*-independent manner. This data is congruous with the established role of *sex-1* in promoting the hermaphrodite sex determination pathway, and the *xol-1*-independent contribution of *sex-1* activity towards hermaphrodite sex determination could be attributed to its repressive effect on *her-1a*. Within the sex determination pathway (Fig. 1A), *her-1* and *tra-1* are the only two genes that are known to be regulated at the level of transcripts [8], [38], [39].

**Fig. 5:**
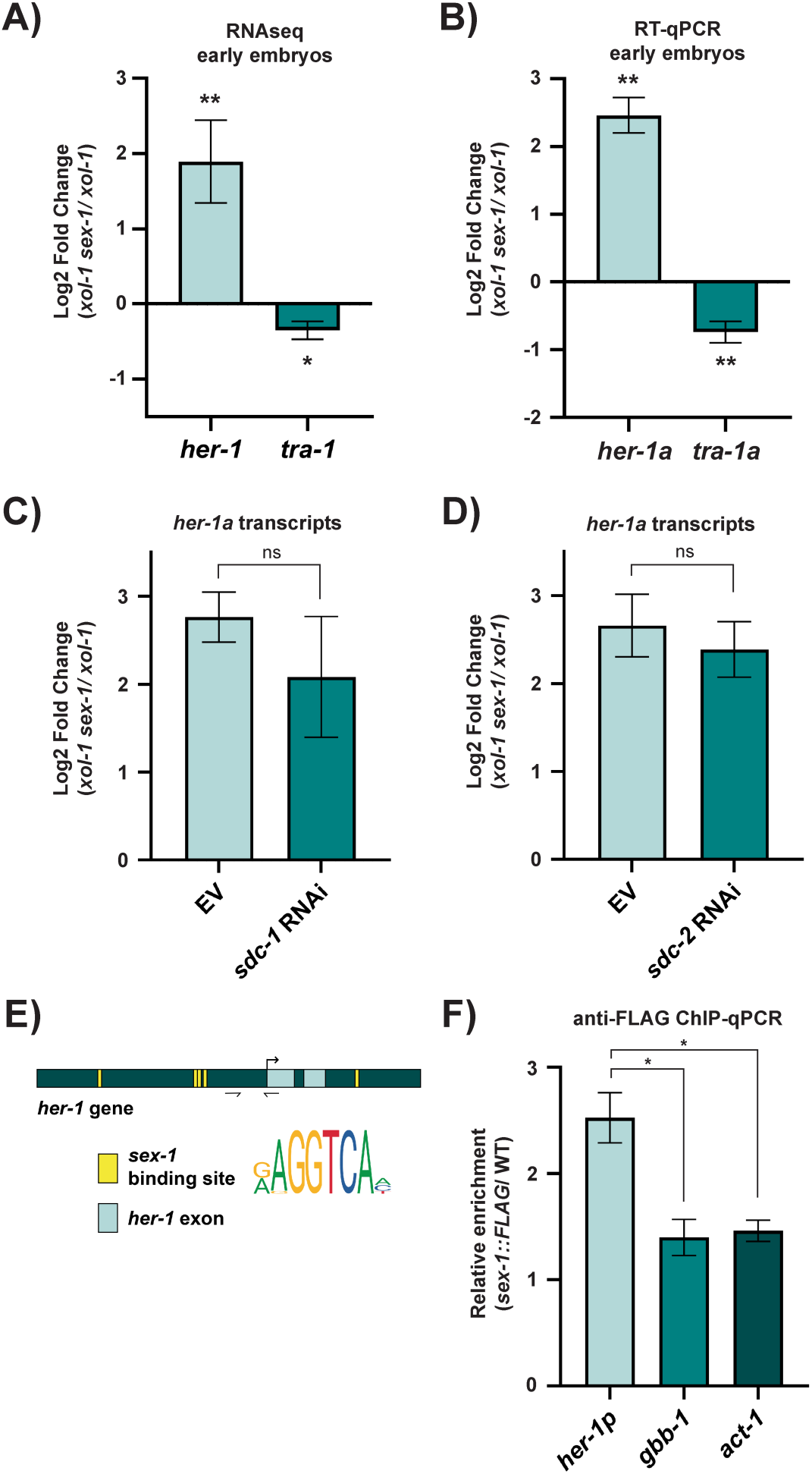
*her-1* is a direct transcriptional target of *sex-1*. (A) Log2 fold change in *her-1* (p = 0.003) and *tra-1* (p = 0.023) in *xol-1 sex-1*/ *xol-1* early embryo RNA-seq datasets. Error bars indicate IfcSE. Statistical significance indicated are the adjusted p-values reported by DESeq2 (wald test with Benjamini-Hochberg correction). n = 3. (B) Log2 fold change in *her-1a* (p = 0.002) and *tra-1a* (p = 0.005) transcripts in *xol-1 sex-1*/ *xol-1* early embryos validated using RT-qPCR. Error bars indicate SEM. P-values obtained from Welch’s unpaired t-test with unequal variance. n = 3. (C-D) Log2 fold change in *her-1a* transcripts upon treatment with empty vector (EV) and *sdc-2* RNAi (p = 0.218) or *sdc-1* RNAi (p = 0.329). Error bars indicate SEM. P-values obtained from Welch’s paired t-test with unequal variance. n = 3. (E) Schematic of *her-1* gene depicting predicted SEX-1 binding sites identified using MEME Suite tool FIMO (Find Individual MOTIF Occurences) [40]. ChIP primers are indicated using arrows. (F) ChIP-qPCR against SEX-1:FLAG shows relative enrichment of *her-1* promoter fragment. Error bars indicate SEM. P-values obtained from Welch’s paired t-test with unequal variance. n = 3. Asterisks indicate level of statistical significance (* p < 0.05, ** p < 0.01, *** p < 0.005, ns not significant).

There are two possible mechanisms through which *sex-1* may be regulating *her-1*. The first is direct transcriptional repression through SEX-1 binding to regulatory elements in the *her-1* promoter. The second is a mechanism where *sex-1* may be acting via the SDC proteins to enable their repression of *her-1*. From the canonical *sex-1* pathway (Fig. 1A), we know that *xol-1* represses the activity of the *sdc* genes, which would otherwise lead to the transcriptional repression of *her-1*. The SDC proteins associate with each other to form a complex and carry out this repression, with SDC-2 and SDC-3 playing the key role in initiating the assembly of the SDC complex onto the *her-1* gene. SDC-1, SDC-2 and SDC-3 together promote this inhibition of *her-1*, with SDC-1 and SDC-2 playing a more prominent role [9]. The presence of functional *sdc-3* gene is required for complete repression of *her-1*, but it is not the rate-limiting protein in this complex [9].

This raises the question of whether *sex-1* acts through and/or in conjunction with the SDC proteins to act upon the *her-1* gene. To test this, we subjected our WT, *xol-1* and *xol-1 sex-1* strains to RNAi treatment against *sdc-1* and *sdc-2* to deplete these proteins. We started RNAi treatment at L1 larval stages, so that embryos formed from the adult germline are depleted in *sdc-1* and *sdc-2*. We measured functional RNAi efficacy using *her-1a* upregulation caused by the repression of *sdc* genes as a readout (Fig. S3A). We found that *her-1a* was upregulated following RNAi treatment in WT, *xol-1* and *xol-1 sex-1* strains, and that there were no statistically significant differences in RNAi efficacy between the three strains (Fig. S3A). According to the model described above, if *sex-1* requires the presence of SDC-1 and SDC-2 proteins for its *xol-1*-independent regulation of *her-1*, the *her-1* upregulation seen in *xol-1 sex- 1*/ *xol-1* should be diminished when *sdc-1* or *sdc-2* is depleted in both *xol-1 sex-1* and *xol-1* strains. However, if *sex-1* does not require the activity of the SDC complex, then there should be no statistically significant difference in *her-1* upregulation after RNAi. We saw no statistically significant difference in *her-1* upregulation in *xol-1 sex-1*/ *xol-1* between empty vector (EV) and *sdc-1* RNAi (Fig. 5C). Both the EV and *sdc-1* RNAi condition showed *her-1* upregulation of ∼2.7 log2 fold change. Similarly, the *sdc-2* RNAi treatment showed no statistically significant difference in *her-1* upregulation, which was about ∼2.6 log2 fold change (Fig. 5D). This data suggests that *sex-1* is regulating *her-1* through a mechanism that is independent of *xol-1*, *sdc-1* and *sdc-2*. Since *xol-1* and the SDC proteins are the only characterized upstream regulators of *her-1*, this data raises the possibility of *her-1* being a direct target of *sex-1*.

To determine the precise mechanism of how *sex-1* may be regulating *her-1*, we first examined the sequence of the *her-1* promoter. SEX-1 is known to be capable of binding to NHR consensus sites to repress transcription of its target gene *xol-1* [1]. We therefore looked for conserved *sex-1* binding sites in the *her-1* promoter. Using tools from the MEME Suite for MOTIF analysis, we detected five putative *sex-1* binding sites near the *her-1* promoter region (Fig. 5E). Four of these *sex-1* binding sites are in the promoter sequence of *her-1*, and one of the sites is in the second intron of the *her-1* gene. *her-1a*, the *her-1* splice isoform that regulates male development, includes the first two exons of the *her-1* gene. The presence of *sex-1* binding consensus sequences in the *her-1* promoter and gene body supports a mechanism where *sex-1* directly regulates the expression of *her-1a* transcripts.

To confirm that SEX-1 is capable of binding to these sites in the *her-1* promoter, we used the SEX-1::FLAG strain to pull down DNA sequences associated with SEX-1 in early embryos. The ChIP-qPCR revealed that SEX-1 binds to the *her-1* promoter (Fig. 5F). We saw an enrichment of *her-1* promoter fragments in SEX-1::FLAG compared to the WT control. As control, we also performed ChIP-qPCR on the *act-1* gene, which is present on the same chromosome as *her- 1* (chromosome V), and on the *gbb-1* gene, which is present on the X chromosome. This enrichment of *her-1* promoter fragments over control fragments suggests that SEX-1 binds directly to the *her-1* promoter to repress its transcription, in a mechanism similar to its repression of the *xol-1* gene.

### *sex-1* regulates X chromosome gene expression

In addition to its effects in the regulation of sex determination, previous studies from our lab and others using the *xol-1* male rescue assay have suggested that *sex-1* also has a *xol-1*- independent role in promoting dosage compensation [2], [33]. While *sex-1* may be regulating sex determination directly through *her-1*, dosage compensation is independently mediated by the SDC proteins parallel to *her-1*. This implies the presence of additional *sex-1* targets that specifically regulate the process of dosage compensation.

To characterize this function of *sex-1*, we first quantified the degree of dosage compensation disruption in *xol-1 sex-1* double mutants. We performed RNA sequencing on WT, *xol-1*, and *xol-1 sex-1* synchronized L1, as the establishment of dosage compensated X chromosomes is first measurable by RNA-seq at the L1 larval stage [27], [41]. To measure defects at a later developmental stage, we also examined L3 larvae. At this stage, the germline is relatively undeveloped allowing analysis of dosage compensation in predominantly somatic tissue [42]. We examined *xol-1*/ WT and *xol-1 sex-1/ xol-1* datasets at the L1 and L3 larval stages, and observed that there were not any large changes in global expression for X-linked genes (Fig. 6A-B, Fig. S4A-B). However, further analysis showed that specific subsets of X-linked genes were significantly more affected by the loss of *xol-1* and *sex-1*. In Fig. 6C-D, we used datasets published in Trombley at el. (2024) [43] to categorize X-linked genes into those that are sensitive to the loss of DPY-27 (302 genes), a component of the DCC, displaying at least a 2-fold upregulation in gene expression in its absence, and those that do not fall into this category were considered to be not sensitive to the loss of DPY-27 function (2018 genes) at the L3 stage. We found that DPY-27 sensitive X-linked genes are significantly repressed due the loss of *xol-1* (Fig. 6C), and that these same genes are significantly upregulated due to the loss of *xol-1*-independent sex*-1* function (Fig. 6D). Using a similar analysis, we examined the effect of *xol-1* and *sex-1* on genes that are sensitive to the loss of DPY-21 (729 genes), another component of the DCC that is also responsible for the deposition of H4K20me1 on the X chromosome. We found a similar pattern where DPY-21 sensitive X-linked genes were preferentially repressed by the loss of *xol-1* and upregulated due to the loss of *xol-1*- independent *sex-1* function (Fig. S4C-D). We examined the overlap of X-genes that are sensitive to the loss of DPY-27, DPY-21 and SEX-1, and found that while only a few genes were common between all of these dosage compensation mutants, 40% of SEX-1 sensitive genes (106 genes) were also sensitive to the loss of DPY-21 and/or DPY-27 (Fig. S4E). The upregulation of DPY-27 sensitive X-genes in the *xol-1 sex-1/ xol-1* dataset is 0.23 (Fig. 6D), and the upregulation of DPY-21 sensitive X-genes is 0.17 (Fig. S4D). This magnitude of change is in line with X chromosome derepression seen in other mild dosage compensation mutants, such as *dpy-27* RNAi, *set-4* mutant and some H3K9me mutants [26], [41]. The X derepression of DCC-sensitive genes in the *xol-1 sex-1/xol-1* comparison confirms that *sex- 1* mediates dosage compensation independent of *xol-1*, through a mechanism that targets a specific subset of X-linked genes and reinforces DCC-mediated dosage compensation.

**Fig. 6:**
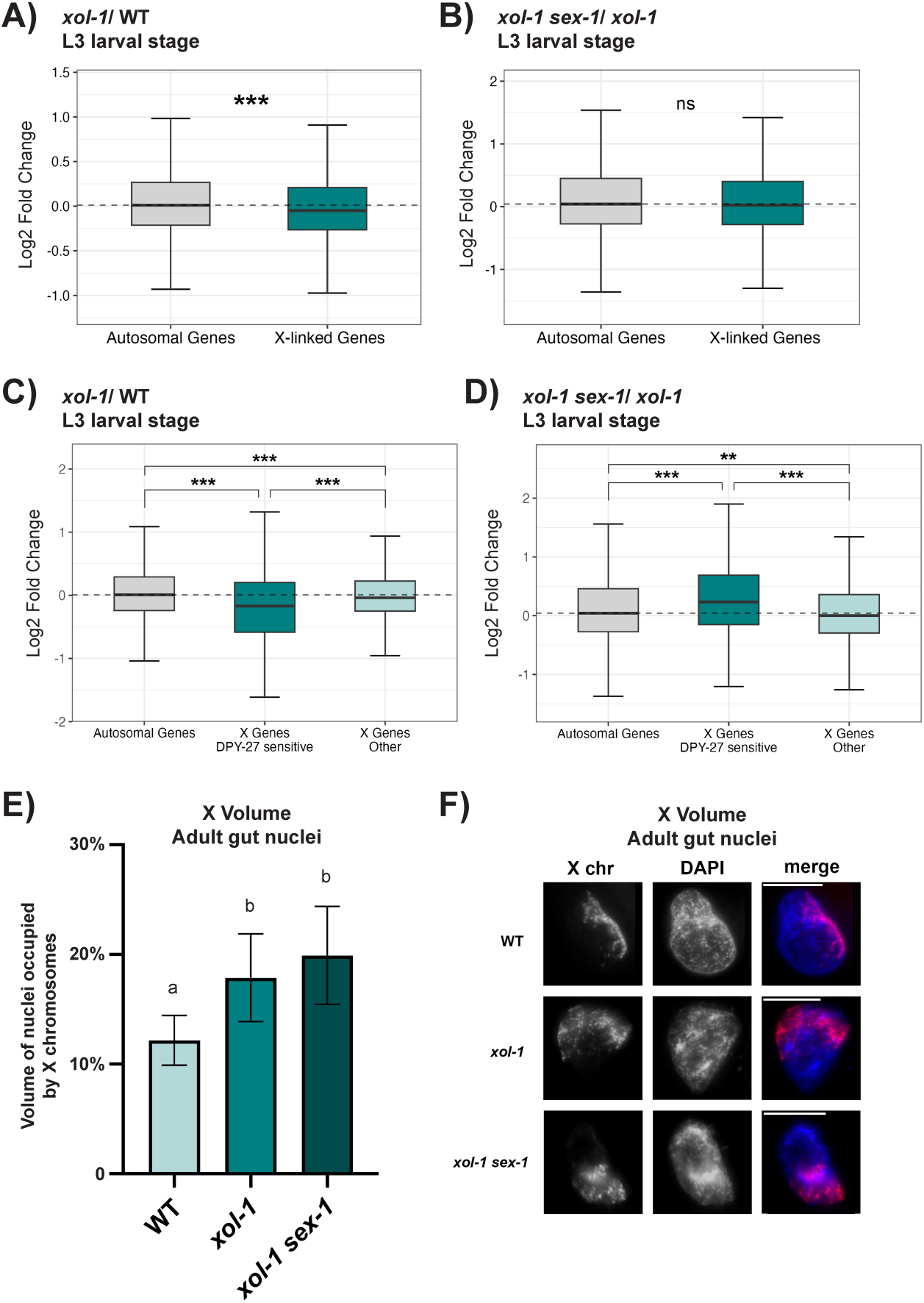
*sex-1* regulates X chromosome gene expression. (A-B) Boxplots showing the distribution of log2 fold change for genes on the autosomes and X chromosomes in (A) *xol-1*/ WT (median log2 fold change (X - A) = -0.06) and (B) *xol-1 sex-1/ xol-1* (median log2 fold change (X - A) = -0.01) at the L3 larval stage. n = 4. Statistical significance was obtained using Wilcoxon rank-sum test. Asterisks indicate level of statistical significance (* p < 0.05, ** p < 0.005, *** p < 0.0005, ns not significant). (C-D) Boxplots showing the distribution of log2 fold change for genes on the autosomes, X-genes sensitive to the loss of DPY-27 function, and X-genes that are not sensitive to the loss of DPY-27 function, in (C) *xol-1*/ WT and (D) *xol-1 sex-1*/ *xol-1*. DPY-27 sensitive X-genes were defined by genes that demonstrate log2 fold change > 1 due to the loss of DPY-27 function. DPY-27 sensitive genes were obtained from analysis of datasets published in Trombley et al. 2024 [43]. Statistical significance was obtained using Wilcoxon rank-sum test. Asterisks indicate level of statistical significance (* p < 0.05, ** p < 0.005, *** p < 0.0005, ns not significant). (E) X volume quantification from IF imaging against DPY-27 to mark the X chromosomes in WT, *xol-1* and *xol-1 sex-1* strains. Y-axis represents the proportion of total nuclear volume occupied by the X chromosomes. Error bars indicate SEM. P-values obtained from Welch’s unpaired t-test with unequal variance. P-values reported in Table S1. 20 distinct nuclei were quantified from at least 10 biological replicates. (F) Representative IF images quantified in (A). Scale bars represent 10 µm.

One of the classic phenotypes of disrupted dosage compensation is decondensation of the X chromosomes. X chromosomes are maintained inside the nuclei in a condensed form through the action of the DCC [44]. When dosage compensation is disrupted, the X chromosomes appear physically decondensed and occupy a greater volume within the nucleus. To measure this phenotype in our mutants, we performed immunofluorescence (IF) imaging using anti-DPY-27 antibodies. The DCC-component DPY-27 associates with the X chromosomes in interphase nuclei. Using this IF assay, X chromosomes in the *xol-1 sex-1* strain appear to occupy a much greater volume of the nuclei than in WT (Fig. 6E-F). However, the *xol-1* mutant worms also have decondensed X chromosomes compared to WT (Fig. 6E-F). The X volume in *xol-1 sex-1* double mutants was not statistically significantly different from *xol-1* mutant alone, suggesting that the X chromosome decondensation is the result of *xol-1*-based misregulation in the dosage compensation pathway. In both the *xol-1* and *xol-1 sex-1* strains, the X chromosomes are occupying ∼17-19% of the nuclear volume. This suggests that the non-canonical *xol-1*-indepedent *sex-1* function in dosage compensation does not contribute towards the physical condensation of the X chromosome, which is in line with the sequencing data that demonstrates a lack of global X derepression in the absence of *sex-1* (Fig 6B).

### *sex-1* regulates the deposition of H4K20me1

To determine the mechanism of *sex-1*’s regulation of dosage compensation, we looked for misregulation of genes that are known to be involved in promoting dosage compensation. We found no differential expression of the known DCC components *dpy-27*, *dpy-26*, *dpy-28*, *dpy-30*, *capg-1*, *mix-1*, *sdc-1*, *sdc-2*, and *sdc-3* using RT-qPCR. However, we found *dpy-21*, one of the components of the DCC, to be statistically significantly downregulated in *xol-1 sex-1* mutant early embryos compared to *xol-1* (Fig. 7A). *dpy-21* is also upregulated in *xol-1*/ WT, demonstrating antagonistic regulation by the *xol-1* and *sex-1* pathways similar to what was seen in the case of *her-1a*. *dpy-21* is responsible for the enrichment of the repressive H4K20me1 histone modification on the X chromosomes [23], [24], [25], [41]. Mutations in *dpy-21* lead to X chromosome derepression that is measurable starting at the L1 larval stage [41]. The enrichment of H4K20me1 on the X chromosomes is also lost when *dpy-21* is mutated [24]. Therefore, the downregulation of *dpy-21* could explain the dosage compensation defects seen in the *xol-1 sex-1* mutants. To test this, we first measured whether H4K20me1 was enriched on the X chromosomes in the *xol-1 sex-1* mutant adult gut nuclei. We performed IF imaging using H4K20me1 antibody to measure the intensity of the histone mark on the X chromosomes normalized to the entire nucleus (see Methods). We used CAPG-1 antibodies, one of the components of the condensin I^DC^, as a co-stain to mark the X chromosomes. Using this assay, we found that H4K20me1 was statistically significantly less enriched on the X chromosomes in *xol-1 sex-1* mutants compared to *xol-1* mutants (Fig. 7B-C). Unexpectedly, we found that *xol-1* mutants had much higher H4K20me1 enrichment compared to WT, and that *xol-1 sex-1* mutants rescued this aberrantly high enrichment to slightly lower than WT levels.

**Fig. 7:**
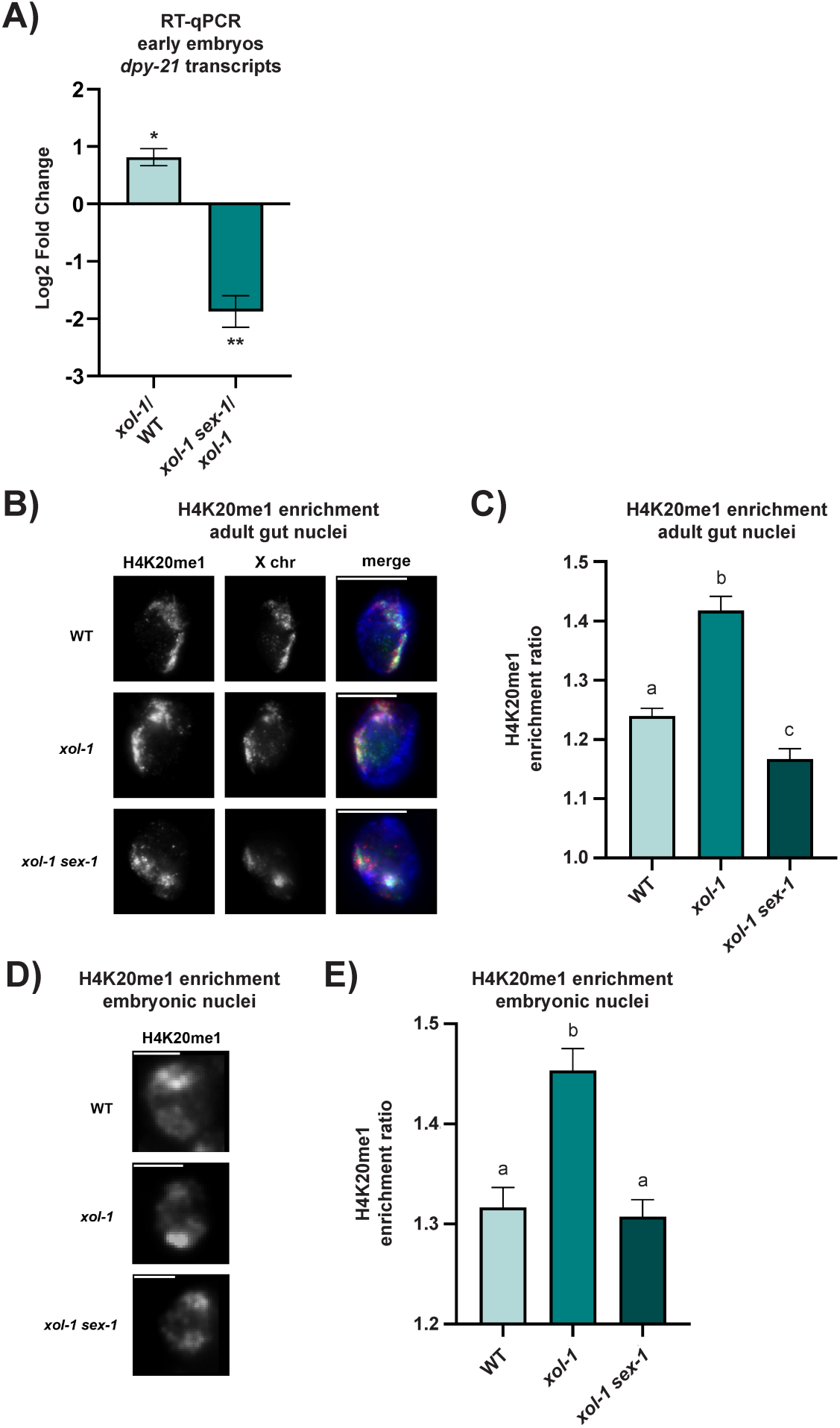
*sex-1* regulates the deposition of H4K20me1. (A) Log2 fold change in *dpy-21* transcripts using RT-qPCR in *xol-1*/ WT (p = 0.019) and *xol-1 sex-1*/ *xol-1* (p = 0.007). Error bars indicate SEM. P-values obtained from Welch’s unpaired t-test. n = 3. Asterisks indicate level of statistical significance (* p < 0.05, ** p < 0.01, *** p < 0.005, ns not significant). (B) Representative images from immunofluorescence staining against H4K20me1 (red) and the X chromosome (CAPG-1, green) in adult gut nuclei in WT, *xol-1* and *xol-1 sex-1*. Scale bars represent 10 µm. (C) Quantification of H4K20me1 intensity in images from (B), scoring H4K20me1 enrichment on the X chromosomes over total nuclei. Error bars indicate SEM. P-values obtained from Welch’s unpaired t-test with unequal variance. P-values reported in Table S1. At least 18 nuclei were quantified per genotype from 10 biological replicates. (D) Representative images from immunofluorescence staining against H4K20me1 in post-mitotic embryonic nuclei in WT, *xol-1* and *xol-1 sex-1*. Scale bars represent 2 µm. (E) Quantification of H4K20me1 intensity in images from (D) scoring H4K20me1 enrichment using line-scan analysis over visibly enriched nuclear loci over visibly non-enriched loci. Error bars indicate SEM. P-values obtained from Welch’s unpaired t-test with unequal variance. P-values reported in Table S1. n = 50.

Since *sex-1* is only expressed in embryonic stages, it is possible that disruptions in dosage compensation and H4K20me1 enrichment during embryogenesis are compensated for later in development. Therefore, we tested the enrichment of H4K20me1 on the X chromosomes in embryonic stages. H4K20me1 is first enriched on the X in >200 cells, with consistent enrichment across nuclei visible in 300- to 350-cell embryos. Unlike the large nuclei of the adult gut cells, the nuclei in these late-stage embryos are very small. To measure the H4K20me1 enrichment in these embryos, we performed immunofluorescence staining with H4K20me1 antibodies. We then employed line-scan analysis to measure the ratio of H4K20me1 enrichment between the X chromosomes and the rest of the nuclei (see Methods) (Fig. 7D-E). We found a similar pattern of enrichment as adult gut cells, where the X chromosomes in *xol-1* mutants had more H4K20me1 enrichment compared to WT, and the addition of a *sex-1* mutation in *xol-1 sex-1* mutants rescued this back to WT levels. Prior studies from our lab and others have demonstrated that *dpy-21* null mutants do not have any measurable H4K20me1 enrichment on the X chromosomes [23], [24], [25], but despite this they do not exhibit defects in developmental viability comparable to those seen in *xol-1 sex-1* mutants [24], [43]. Our data demonstrates that *xol-1 sex-1* mutants exhibit high embryonic lethality, larval lethality, and extremely low fecundity when compared to WT (Fig. 2B-D) but minimal disruption in H4K20me1, indicating that misregulation of dosage compensation via H4K20me1-mediated repression is not likely to be the driver of developmental lethality in these worms.

## Discussion

Our results suggest that *sex-1* plays important roles in the regulation of sex determination and dosage compensation downstream of *xol-1*. SEX-1’s roles in embryonic sex determination are likely mediated through direct transcriptional regulation of some of the important components of these pathways such as *her-1* (Fig. 8). This multi-level regulation enables *sex-1* to reinforce the activation of the hermaphrodite sex development pathway and the establishment of dosage compensation in XX embryos (Fig. 8). We also demonstrate that *sex-1* regulates the enrichment of repressive H4K20me1 on the X chromosomes through transcriptional regulation of the H4K20 demethylase *dpy-21* (Fig. 8), but that this function of *sex-1* is likely not the driver of developmental lethality caused by the loss of *sex-1* in a *xol-1* mutant background.

**Figure 8:**
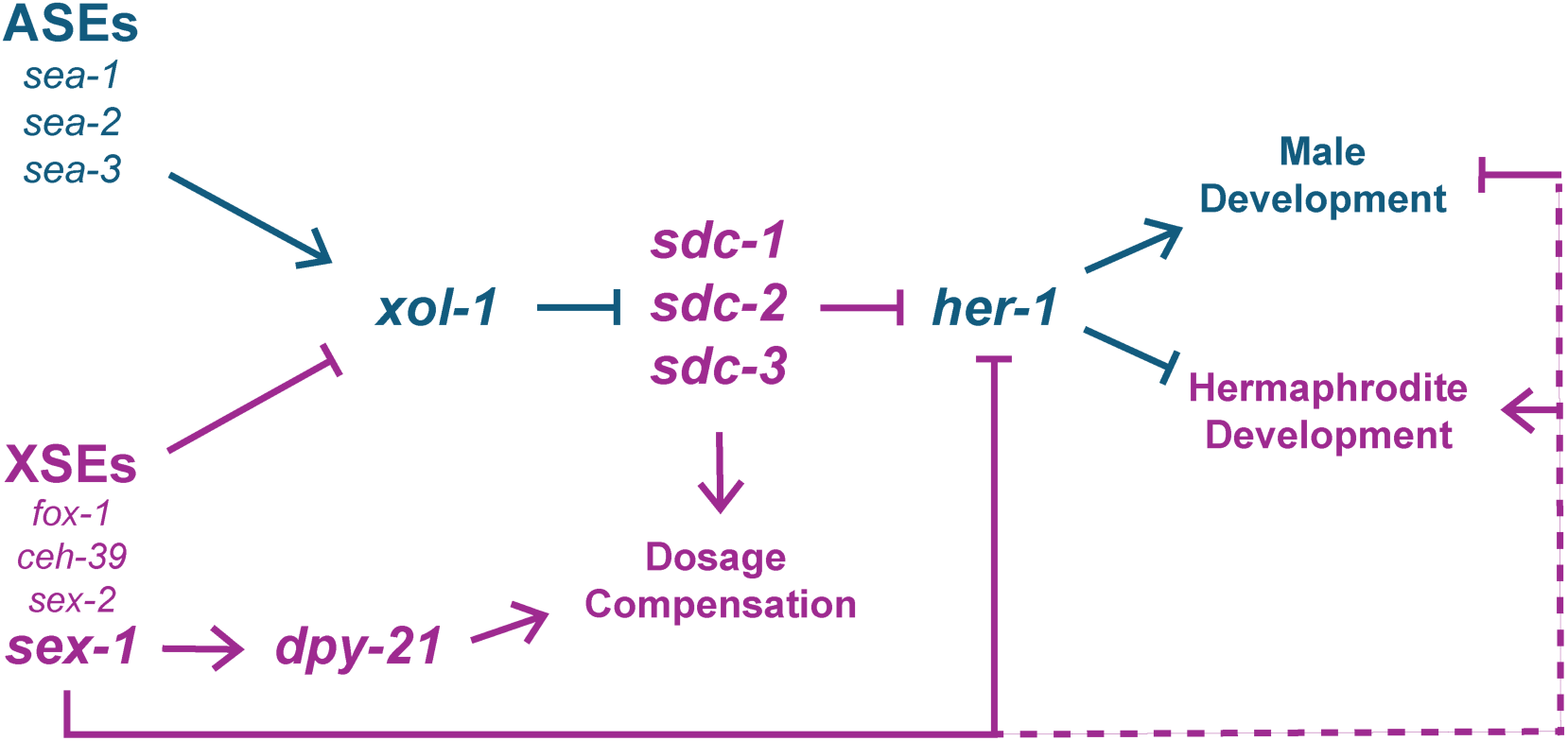
Model for the action of SEX-1 in XX hermaphrodite embryos.

### Shared downstream targets of *xol-1* and *xol-1*-independent *sex-1* regulation

Multiple lines of evidence demonstrates that *sex-1* regulates the downstream targets of *xol- 1,* antagonizing the pathways activated by *xol-1*. These include both the transcriptional targets that are associated with *xol-1*-mediated control of developmental timing, and *xol-1*- mediated regulation of sex-biased transcriptional pathways. Furthermore, there is a significant overlap in genes that are significantly differentially expressed by the two regulators. *her-1*, a canonical downstream target of *xol-1*, was significantly upregulated by the loss of *sex-1* (Fig. 5A-B). SEX-1 was also shown to be able to bind to the *her-1* promoter (Fig. 5F). This validates *her-1* as a target of both *xol-1* and *sex-1*, and this pathway likely contributes to the regulation of the sex-biased transcriptional programs by *sex-1*. In sum, this suggests a model where *sex-1* antagonizes some *xol-1*-mediated pathways through direct transcriptional regulation of *xol-1* targets.

In a previous study by our lab, *met-2* was implicated as a downstream target of the *xol-1* pathway in hermaphrodites. *met-2* was shown to regulate the timing of embryonic development and the onset of dosage compensation [27]. However, *met-2* was not statistically significantly changed between *xol-1 sex-1* and *xol-1* mutants (Fig. S1C). This implies that *met-2* is a target of *xol-1*, but not *sex-1*. *sex-1* likely regulates downstream targets other than *met-2* that ultimately regulate the process of embryonic development. Since *met- 2* is not misregulated, this *sex-1*-mediated control could be through direct regulation of targets downstream of *met-2*. Another possibility is that *sex-1* may regulate parallel pathways with independent targets of its own that are not downstream of *xol-1*. Though most differentially expressed genes in *xol-1 sex-1*/ *xol-1* are also misregulated by *xol-1*/ WT, there are a few significant genes that are only affected by loss of *sex-1* and not *xol-1*. These downstream targets could mediate parallel pathways that feed into control over embryonic development.

### *her-1*-mediated *sex-1* regulation of sex determination

Evidence from our study suggests that *her-1* is a direct transcriptional target whose expression is repressed by SEX-1 in hermaphrodite embryos. An important question that arises from this result is whether transcriptional control of the *her-1* promoter can explain all of *sex-1*’s regulation of sex determination in *C. elegans*. This appears to be a possibility as *her-1* is the most potent known regulator of male sex-specific embryonic development. Very low-levels of *her-1* expression is sufficient to induce male-specific cellular differentiation in embryos [7], [8]. While *xol-1* expression is important during early embryogenesis, tight control over *her-1* regulation is necessary all throughout *C. elegans* development as continued *her-1* expression during larval stages is necessary to complete male-specific somatic development in XO animals [7], [8], [45].

However, evidence from looking at the differences between hermaphrodite- and male- biased transcriptional gene sets suggests that *her-1* is likely not the only target of *sex-1* within the sex determination pathways. If *her-1* was the primary target of *sex-1* beyond its *xol-1* regulation, we would expect a greater degree of change in the male-biased transcriptional program upon the loss of *xol-1*-independent *sex-1* function in the *xol-1 sex-1/ xol-1* dataset (Fig. 4D). The male-biased pathways are indeed upregulated by the loss of *sex-1*, but they remain suppressed compared to the WT condition (Fig. 4D). In contrast, the hermaphrodite sex-biased pathways are significantly more affected by the loss of *sex-1* (Fig. 4C). This difference in magnitude of change between the male- and hermaphrodite-biased genes suggests that *sex-1* likely has more targets than *her-1*, and that many of these targets may act specifically on hermaphrodite sex determination.

### Multi-level feedback in transcriptional networks governing sex determination

The direct regulation by *sex-1* on the downstream targets of *xol-1* represents a multi-level regulatory paradigm where the transcriptional output of the sex determination and dosage compensation is stabilized at several levels by the upstream regulator *sex-1*. Multi-level regulation in the form of feedback loops are also seen by other components of the *C. elegans* sex determination pathway. For example, *tra-1* exerts similar multi-level feedback by repressing the transcription of its male development-promoting upstream regulators *fem-3, xol-1* and *sup-26* [39], [46]. Another feedback loop within *C. elegans* dosage compensation is the regulation of gene expression of XSEs *sex-1, fox-1* and *ceh-39* that are all dosage compensated by the DCC [2]. In this feedback loop, *sex-1* is itself transcriptionally repressed through the process of dosage compensation, and therefore *sex-1* regulates its own activity through a negative feedback loop. However, neither of these feedback loops are similar to the action of *sex-1* on multiple levels of its downstream pathway.

### Mechanism of action of NHR SEX-1

*C. elegans* contains many more NHRs than most organisms, representing a unique evolutionary expansion of this superfamily [47]. Though many *C. elegans* NHRs do not have any close homologs in other organisms, SEX-1 has close homologs in mammalian Rev-erb and *Drosophila* E78A [28]. There is little functional conservation between Rev-erb and E78A beyond their important roles in development. Rev-erb is involved in the function of the circadian clock in mammals, and is thought to have uncharacterized roles in mammalian embryonic development [48], [49], [50]. Drosophila E78A regulates lipid metabolism, and has important roles in regulating molting and larval development [51]. In the initial studies characterizing SEX-1’s mechanism of action, it was thought to be similar to Rev-erb which was at the time classified as an orphan NHR [28]. Rev-erb is also a transcriptional repressor, similar to the known roles of *sex-1*. However, more recent studies have revealed heme to be an important ligand regulating Rev-erb activity [52]. The high divergence between the *sex-1* LDB and LBDs from NHRs from other organisms suggests that either *sex-1* is an orphan NHR, or that it may be potentially able to bind uncharacterized *C. elegans*-specific ligands [28].

NHRs also often interact with binding partners such as coactivator and corepressor complex proteins to exert their transcriptional control over target genes [53]. Since *sex-1* represses *her-1*, and *her-1* is also the primary target of the SDC proteins that are involved in both sex determination and dosage compensation, a reasonable hypothesis arose which stated that *sex-1* may interact with the SDC proteins to repress *her-1*. However, RNAi treatment against the rate-limiting *sdc* genes *sdc-1* and *sdc-2* demonstrates that *her-1* remains upregulated in the absence of *sex-1* function regardless of the presence of the SDC proteins (Fig. 5C-D). This data leads us to conclude that *sex-1* acts independently from the *sdc* proteins. Therefore, *sex-1* acts on shared targets of the *xol-1* and *sdc* complex-mediated pathway, independently from both these components.

### *sex-1* acts on specific X-linked genes to promote dosage compensation

Prior studies exploring the downstream targets of *sex-1* suggested that the developmental lethality observed in *xol-1 sex-1* double mutants is a consequence of disruption in dosage compensation [2]. However, we found no statistically significant increase in X volume in *xol- 1 sex-1* double mutants compared to *xol-1* mutants alone (Fig. 6E-F). Additionally, we observed no significant global changes in X chromosome gene expression in *xol-1 sex-1*/ *xol- 1* L1 and L3 larval datasets (Fig. 6A-B, Fig. S4A-B). However, closer analysis of these datasets revealed that specific subsets of X-linked genes that are sensitive to the loss of DPY-27 and/or DPY-21, show statistically significant derepression in the *xol-1 sex-1*/ *xol-1* dataset (Fig. 6C-D, Fig. S4C-D). This suggests that *sex-1* likely reinforces DCC-mediated repression of a subset of specific target X-linked genes. Loss of function *sex-1* mutations have been frequently used as “sensitizer” mutations that predispose worms towards developing dosage compensation defects upon partial RNAi against genes that mediate dosage compensation [26], [33], [54]. This mechanism explains why the loss of *sex-1*-mediated reinforcement of dosage compensation may sensitize worms to further disruptions in the dosage compensation pathway.

Evidence from our study suggests that *sex-1*, either directly or indirectly, promotes the expression of *dpy-21* in embryos in a *xol-1*-independent manner (Fig. 7A). The loss of *sex-1* in a *xol-1* mutant background also leads to a decrease in the enrichment of repressive H4K20me1 on the X chromosomes compared to *xol-1* mutant alone, but is only slightly decreased when compared to WT (Fig. 7B-E). This suggests that while *sex-1* may be promoting dosage compensation through the deposition of H4K20me1 in XX embryos, this pathway is likely not responsible for the significant embryonic and larval lethality observed in *xol-1 sex-1* double mutants. Characterization of *dpy-21* null mutants from prior studies have shown that *dpy-21* mutants do not exhibit any significant embryonic lethality [43], even though the disruption of dosage compensation in terms of global derepression of X-linked transcripts is significantly more severe than the subtle derepression of DCC-sensitive X- linked genes observed in *xol-1 sex-1* mutants [43]. H4K20me1 enrichment is also known to be completely abolished in these *dpy-21* mutants, compared to the slight decrease observed in *xol-1 sex-1* vs WT [24]. Together, these data provide further evidence of the presence of additional novel *sex-1* targets that either promote dosage compensation through an as-yet uncharacterized *dpy-21*-independent mechanism, or directly regulate pathways involved in embryonic and/or larval development. Misregulation of these targets may lead to lethality through a mechanism independent from dosage compensation.

### Effect of *xol-1* on dosage compensation beyond embryogenesis

An earlier study published by our lab, Jash et al. (2024), demonstrated that low-levels of *xol- 1* expression was required for the proper timing of initiation of dosage compensation during early embryogenesis [27]. However, the study concluded that beyond this regulation during the early stages of embryo formation, *xol-1* likely does not have any role in the development of post-embryonic tissues. However, evidence from our current study suggests that loss of *xol-1* results is several subtle phenotypes at the L3 larval stage, as well as in adult gut cells. *xol-1*/ WT RNA-seq suggests that *xol-1* mutants significantly repress specific subsets of X-linked genes i.e. those that are sensitive to the loss of DPY-27 and DPY-21 (Fig. 6C, Fig. S4C). This is in line with its known role during hermaphrodite embryogenesis, where it inhibits the dosage compensation pathway, resulting in earlier loading of DCC components in the absence of *xol-1* [27]. Our study shows that in post-mitotic embryonic nuclei, *xol-1* mutants have significantly higher H4K20me1 enrichment on their X chromosomes compared to WT (Fig. 7D-E). Somewhat surprisingly, significantly higher enrichment of H4K20me1 is also observed in *xol-1* mutant adult gut nuclei (Fig. 7B-C). While the higher enrichment in embryonic nuclei is in line with the role of *xol-1* in inhibiting dosage compensation during embryogenesis, its continued aberrant enrichment in adult tissues suggests that *xol-1* may be active beyond the stages of embryonic development, or that changes mediated by XOL-1 can be propagated in a *xol-1* independent manner. Finally, the volume of X chromosome in *xol-1* mutants appears to be significantly higher than in WT. This phenotype generally indicates a disruption in dosage compensation [26], [33], [43], [44], but the RNA-seq data does not show any global X-derepression. These results are similar to *cec-4* mutants, where the X chromosomes are significantly decondensed, but they exhibit either very minimal or no significant X-derepression [26], [43].

## Materials and Methods

### Worm Maintenance

All strains were maintained on NG agar plates with *E. coli* (OP50) as a food source, using standard methods [55]. Strains include: N2 Bristol strain (wild type); WM458 *xol-1(ne4472)* X; VC1703 *+/szT1 [lon-2(e678)]* I*; sex-1(gk829)/szT1* X; EKM163 *sex-1(gk829)* X, EKM198 *xol-1(ne4472) sex-1(gk829)* X, EKM193 *sex-1(cld14[sex-1::FLAG])* X. Some strains were provided by the CGC, which is funded by NIH Office of Research Infrastructure Programs (P40 OD010440).

### CRISPR-CasG gene editing

For the injection procedure, young adult worms were immobilized on agar pads, followed by microinjection with injection mix containing Cas9 protein (PNA Bio CP01), ssDNA repair template with FLAG insert, gRNA targeting the *sex-1* C-terminus, and gRNA and repair template targeting the co-CRISPR site *dpy-10* was injected into the distal and/or proximal gonads of gravid young adult *C. elegans* [56]. From the progeny of injected worms, worms containing the roller phenotype due to the *dpy-10* co-CRISPR were isolated and screened by PCR for FLAG insertion. Correct edit was confirmed by sequencing. After the isolation of homozygous lines, the worms were backcrossed three times. FLAG sequence was inserted just before the stop codon at the C-terminus of the *sex-1* gene. gRNA, repair template and PCR primer sequences are listed in Table S2.

### Viability assays

For embryonic viability, young gravid adults were allowed to lay embryos and the number of embryos on each plate was counted. After 24h at 20°C, embryonic lethality was scored based on the number of embryos remaining on the plate. For larval viability, the same procedure was followed as for embryonic viability, and after another 48h at 20°C, viability was scored based on the number of adult worms on the plate. For calculating embryonic viability, the number of hatched eggs was divided by the total number of eggs laid. For calculating larval viability, the number of surviving adults was divided by the total number of hatched larvae. For progeny quantification, synchronized L1 larvae were plated onto NGM plates with OP-50 and allowed to reach L4 stage. L4 worms were moved onto a new plate. Young adult worms were moved every day until they stopped laying eggs. After 72h at 20°C, young adult worms from each plate were counted. For progeny quantification, the cumulative total progeny for each worm was reported. Statistical significance for all three assays was evaluated using Welch’s unpaired t-tests for each comparison of 2 conditions. For embryonic viability, >12 replicates, representing a total of >1100 embryos, were quantified for each condition. For larval viability, >5 replicates, representing a total of >550 larvae, were quantified for each condition. For progeny quantification, the total progeny from 10 worms were counted for each condition.

### RNAi treatment

The RNAi construct for empty vector control and *sdc-1* RNAi were obtained from the Ahringer RNAi library [57]. *sdc-2* RNAi strain was generated by PCR amplification of target sequence from the *C. elegans* genome, followed by restriction enzyme (RE)-mediated insertion into the L4440 RNAi feeding vector. Primer sequences are listed in Table S2. The insert was then sequence verified. Synchronized L1 larvae were seeded onto RNAi plates with the appropriate HT-115 *E. coli* RNAi strains. The worms were allowed to grow at 20°C for 72h. Young adult worms were bleached to obtain early embryos, which were then frozen in TRIzol (Invitrogen catalog number 15596026) at -80°C.

### RNA extraction

Synchronized gravid adult worms were bleached to obtain early embryos. For L1 larvae, synchronized early embryos were incubated in 1x M9 with shaking for 24h, then the newly hatched L1 larvae were incubated in OP50 for 3 hours. Samples were lysed by repeated freeze-thaw cycles using liquid nitrogen. TRIzol-chloroform (Invitrogen catalog number 15596026, Fisher Scientific catalog number BP1145-1) separation of the samples was followed by total RNA extraction using the QIAGEN RNeasy Mini Kit (Qiagen catalog number 74104) with on-column DNase I digestion using RQ1 RNase-Free DNase (Promega catalog number M6101).

### RT-qPCR

cDNA was generated from extracted RNA using random hexamers with SuperScript III First-Strand Synthesis System (Invitrogen catalog number 18080051). RT-qPCR reaction mix was prepared using Power SYBR Green PCR Master Mix (Applied Biosystems catalog number 43- 676-59) with 10 µl SYBR master mix, 0.8 µl of 10 µM primer mix, 2 µl sample cDNA, and 7.2 µl H2O. Samples were run on the Bio-Rad CFX Connect Real-Time System. Log2-fold change was calculated relative to control (*act-1*). Control primers were taken from Hoogewijs et. al. (2008) [58]. Three replicates were analyzed for each genotype. Statistical significance was calculated using 2-tailed unpaired Welch’s t-test with unequal variance. PCR primer sequences are listed in Table S2.

### mRNA-seq analysis

Poly-A enrichment, library prep and next-generation sequencing was carried out in the Advanced Genomics Core at the University of Michigan. RNA was assessed for quality using the TapeStation or Bioanalyzer (Agilent). Samples with RINs (RNA Integrity Numbers) of 8 or greater were subjected to Poly-A enrichment using the NEBNext Poly(A) mRNA Magnetic Isolation Module (NEB catalog number E7490). NEBNext Ultra II Directional RNA Library Prep Kit for Illumina (catalog number E7760L), and NEBNext Multiplex Oligos for Illumina Unique dual (catalog number E6448S) were then used for library prep. The mRNA was fragmented and copied into first strand cDNA using reverse transcriptase and random primers. The 3’ ends of the cDNA were then adenylated and adapters were ligated. The products were purified and enriched by PCR to create the final cDNA library. Final libraries were checked for quality and quantity by Qubit hsDNA (Thermofisher) and LabChip (Perkin Elmer). The samples were pooled and sequenced on the Illumina NovaSeqX 10B paired-end 150bp, according to manufacturer’s recommended protocols. Bcl2fastq2 Conversion Software (Illumina) was used to generate de-multiplexed Fastq files. The reads were trimmed using CutAdapt v2.3 [59]. FastQC v0.11.8 was used to ensure the quality of data [60]. Reads were mapped to the reference genome WBcel235 and read counts were generated using Salmon v1.9.0 [61]. Differential gene expression analysis was performed using DESeq2 v1.42.0 [62]. Downstream analyses were performed using R scripts and packages. Gene set enrichment analysis was performed using GSEA software v4.3.2 [63].

### Immunofluorescence (IF) staining

For adult gut cell IF, 24h post-L4 hermaphrodite worms were dissected in 1x sperm salts (50 mM Pipes pH 7, 25 mM KCl, 1 mM MgSO_4_, 45 mM NaCl, and 2 mM CaCl_2_) and fixed in 4% paraformaldehyde (PFA) on a glass slide for 5min. Slides were then frozen on dry ice. Using a razor blade, the coverslip was removed by flicking it off the slide. The slides were then washed in 1x PBST (PBS with 0.1% Triton X) 3 times for 10 minutes each. 30µL of diluted primary antibody was applied to each slide, and a piece of parafilm was placed over the spot with the antibody to slow evaporation. The slides were incubated in a humid chamber at room temperature overnight. Slides were washed in 1x PBST 3 times for 10 minutes. The slides were then stained with secondary antibodies diluted to 1:100 in 1x PBST at 37°C for 1 hour. Three more washes in 1x PBST were completed after the secondary antibody incubation. In the third wash, DAPI was diluted in 1x PBST. Slides were mounted with Vectashield (Vector Labs catalog number H-1000-10).

For embryo staining, adult worms were first bleached to obtain embryos. After washing in 1x M9, embryos were fixed using finney fix solution (2% v/v paraformaldehyde, 18% v/v methanol, 11.2% v/v witch’s brew (571mM KCl, 71.4mM NaCl, 14.3mM EGTA, 3.5mM spermidine, 1.4mM spermine, 3.5% BME, 107mM PIPES pH 7.5), 1mM EGTA). Pellets were then frozen in -80°C. Samples were thawed and washed 3x for 15mins each in 1x PBST. Samples were incubated for 2 hours at RT with primary antibodies, and then washed 3x 15mins each in 1x PBST. Samples were then incubated in secondary antibody for 1hr at RT, followed by 3x 15min washes in 1x PBST. Slides were then mounted with Vectashield (Vector Labs catalog number H-1000-10).

The following primary antibodies were used for adult gut nuclei imaging: rabbit anti- H4K20me1 (Abcam ab9051) (1:1000), mouse anti-FLAG M2 antibody (Millipore Sigma F1804) (1:1000 dilution), rabbit anti-DPY-27 [17] (1:200 dilution), and goat anti-CAPG-1 [26] (1:200 dilution). The following primary antibodies were used for embryo imaging: rat anti-HTZ-1 (1:100), rabbit anti-H4K20me1 (Abcam ab9051) (1:1000), and mouse anti-FLAG M2 antibody (Millipore Sigma F1804) (1:1000 dilution). Secondary antibodies used were anti-rabbit FITC (711-095-152) (1:100 dilution), anti-rabbit Cy3 (711-165-152) (1:100 dilution), anti-mouse FITC (715-095-150) (1:100 dilution), anti-mouse Cy3 (715-165-150) (1:100 dilution), anti-rat FITC (712-095-153) (1:100 dilution). All secondary antibodies were purchased from Jackson ImmunoResearch Labs.

### Imaging and quantification

Microscopy was performed using an Olympus BX61 microscope and a 60X APO oil immersion objective. Images were taken with a Hamamatsu ORCA-ER High-Resolution Monochrome Cooled CCD (IEEE 1394) camera. 3D images of nuclei were captured using 0.1-0.2 μm Z-stacks. All images shown are 2D projection images of these Z stacks. Quantification was conducted in the Slidebook 5 program (Intelligent Imaging Innovations). To observe signal depletion, exposure times were standardized to wild type. For H4K20me1 intensity quantification in embryonic nuclei, the line-scan function was used on single Z-stacks of isolated nuclei. The peak signal intensity on the line-scan was divided by the lowest signal intensity to obtain the ratio of relative enrichment of H4K20me1 on the X chromosomes. For intensity measurement on adult gut nuclei, the mask function was used to draw segment masks on the X chromosome and DAPI signals. H4K20me1 intensity on both masks were measured. The sum intensity on the X mask was divided by the sum of intensity on the DAPI and X masks to obtain relative X enrichment of H4K20me1. For X volume quantification, the mask function was used to draw segment masks on adult gut nuclei images using the DAPI and X chromosome signals. The volume of each mask was used to calculate the proportion of the nucleus occupied by the X chromosome.

### Chromatin Immunoprecipitation

To generate samples for ChIP, we bleached synchronized adult worm strains to obtain embryos. Embryos were washed three times in 1x M9. Samples were incubated in 2% PFA for 30min for fixation, followed by glycine treatment to halt the reaction. Samples were washed three times in 1x M9 again. The samples were then resuspended in homogenization buffer (50mM HEPES-KOH pH 7.6, 1mM EDTA, 140mM KCl, 0.5% NP-40, 10% glycerol) and frozen in liquid nitrogen at -80°C. Thawed samples were sonicated to shear chromatin using the Covaris M220 focused-ultrasonicator. For the ChIP, 2mg of samples from WT and *sex- 1::FLAG* strains were thawed and incubated with 4µg anti-FLAG antibodies (Millipore Sigma F1804) for 2h at 4°C. Protein G magnetic beads were blocked using 0.1µg/µl BSA for 30min prior incubation with the lysate at 4°C for 30min. The samples were then washed twice in 100mM KCl ChIP Buffer (50mM HEPES-KOH pH 7.6, 1mM EDTA, 100mM KCl, 0.05% NP-40), twice in 1M KCl ChIP Buffer (50mM HEPES-KOH pH 7.6, 1mM EDTA, 1M KCl, 0.05% NP-40), and then twice in 1x IDTE Buffer (catalog number 11-05-01-05). Elution buffer (10mM Tris-HCl pH 8.0, 1% SDS) was used to elute bound proteins and chromatin. 5M NaCl was added to the samples, followed by overnight incubation at 65°C to de-crosslink eluate. Eluate was then treated with RNase A and proteinase K and incubated for 1h at 60°C. DNA fragments were recovered using the Zymo ChIP DNA Clean and Concentrator kit (catalog number D5205).

## Acknowledgements

This work was supported by the National Institute of General Medical Sciences grants number R01 GM13385801 to GC, and R35GM149543 to GC and the National Science Foundation grant number MCB 1923206 to GC. Some *C. elegans* strains used in this study were obtained from the Caenorhabditis Genetics Center (CGC), which is funded by the NIH Office of Research Infrastructure Program (P40 OD010440). The funders had no role in the study design, data collection and analysis, decision to publish, or preparation of the manuscript.

## Data availability Statement

RNA-seq datasets produced by our lab for this study are available at the NCBI GEO database under GSE262626 and GSE278001. Scripts used to process and analyze datasets can be found at https://github.com/eshnaj/jash_sex1_multilevel_reg_paper.

---

**Fig. S1:**
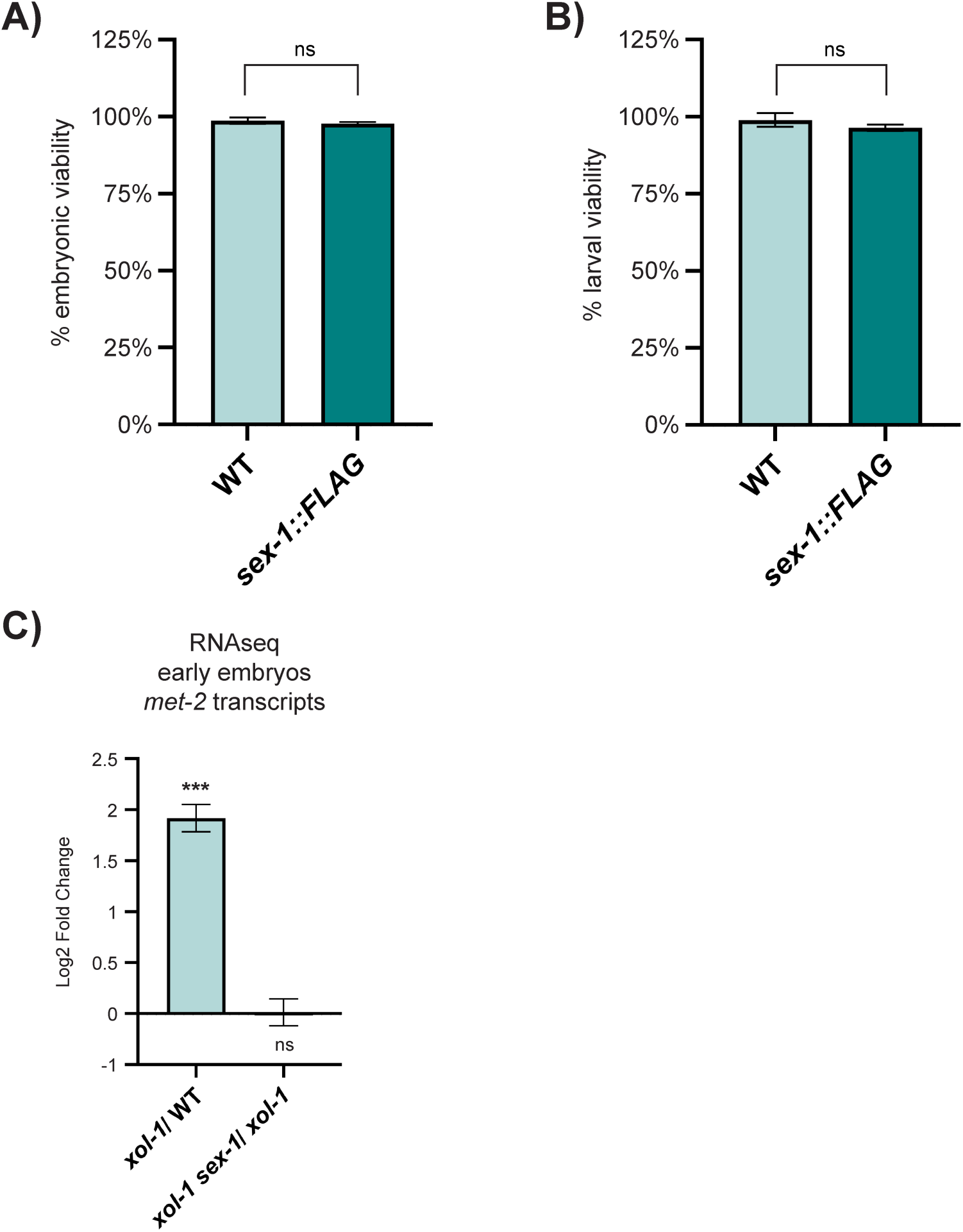
Viability of the sex-1::FLAG strain and *met-2* expression in *xol-1 sex-1* early embryos. (A) Embryonic viability for WT and *sex-1::FLAG* strains (p = 0.694). Error bars indicate SEM. >680 embryos were quantified from at least 9 biological replicates. P-values obtained from unpaired Welch’s test with unequal variance. (B) Larval viability for WT and *sex-1::FLAG* strains (p = 0.092). Error bars indicate SEM. >530 total larvae were quantified from at least 9 biological replicates. P-values obtained from unpaired Welch’s test with unequal variance. (C) Bar chart representing log2 fold change in *met-2* gene expression in *xol-1*/ WT (p = 5.42 x 10^-39^) and *xol-1 sex-1/ xol-1* (p = 0.965) early embryo datasets. Error bars indicate standard error IfcSE. P-values were obtained from adjusted p-values reported by DESeq2 (wald test with Benjamini-Hochberg correction). Asterisks indicate level of statistical significance (* p < 0.05, ** p < 0.01, *** p < 0.005, ns not significant).

**Fig. S2:**
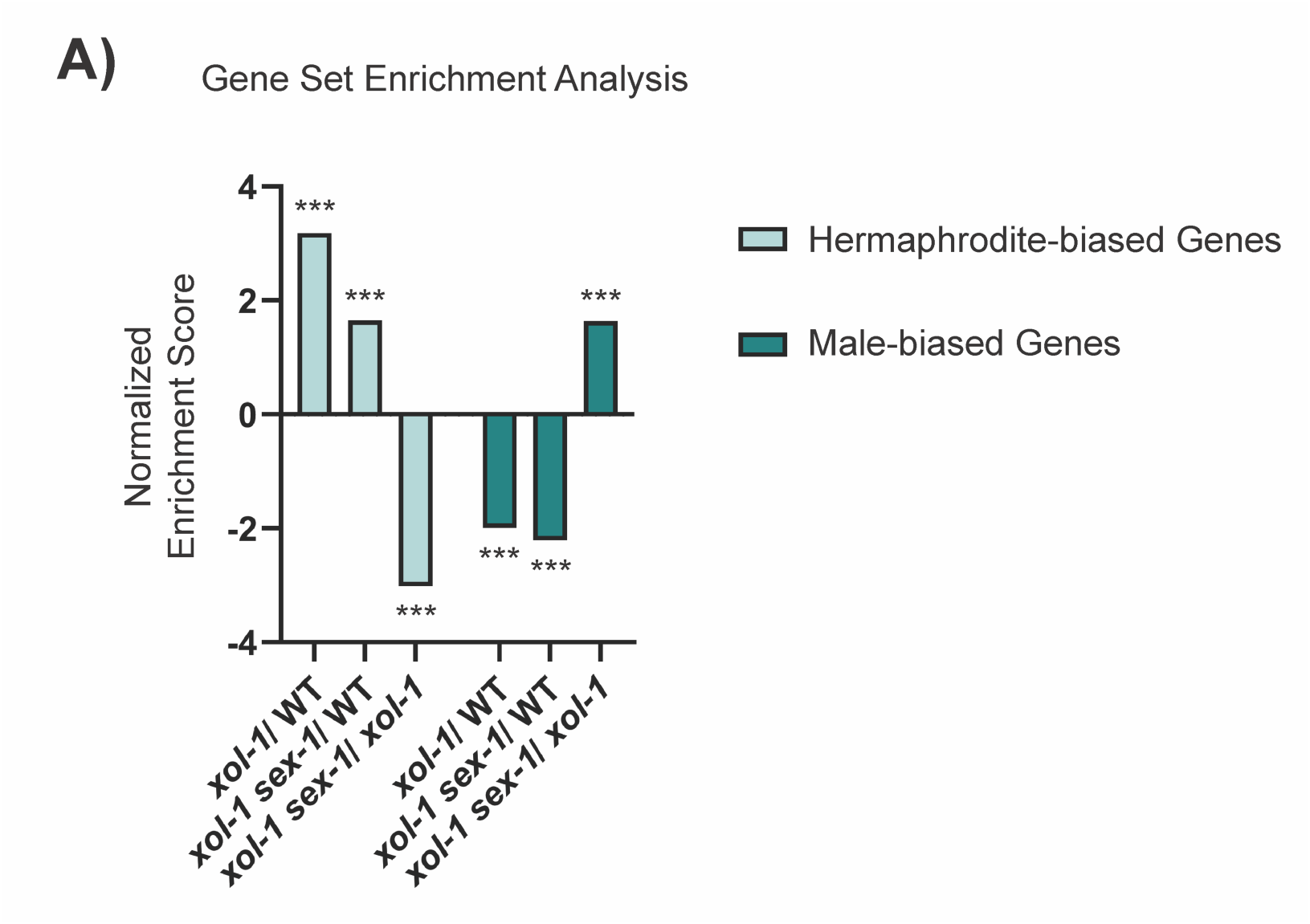
Gene set enrichment analysis. (A) Bar chart with normalized enrichment scores for sex-biased transcripts obtained from Gene Set Enrichment Analysis (GSEA) [27], [63]. Asterisks indicate level of statistical significance (* p < 0.05, ** p < 0.01, *** p < 0.005, ns not significant).

**Fig. S3:**
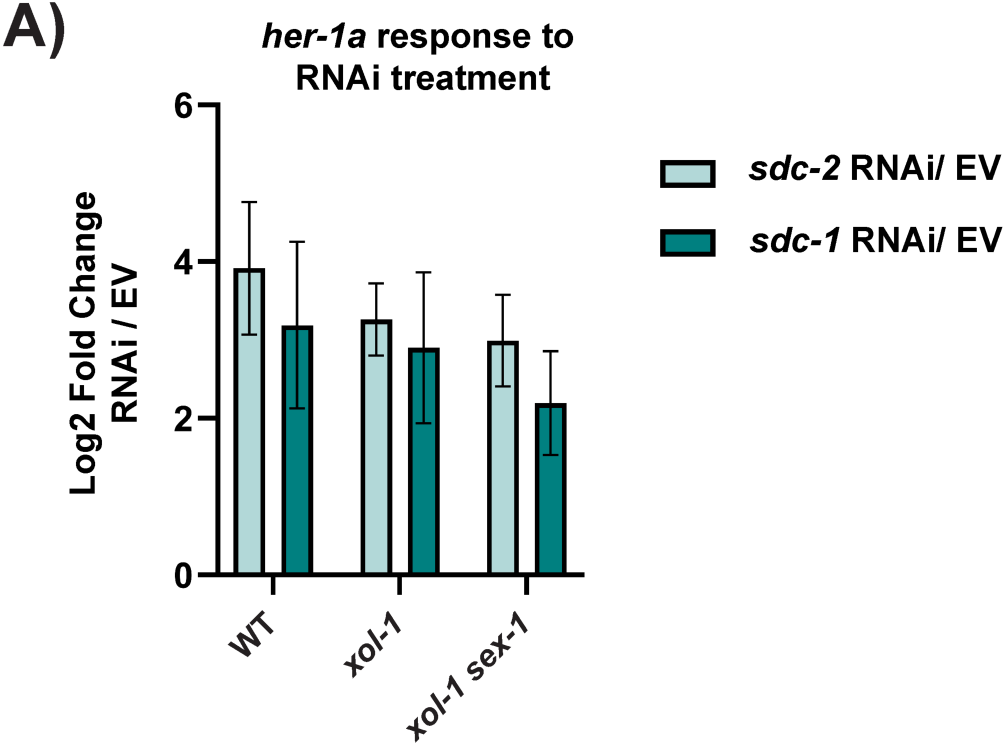
Measuring RNAi efficacy following *sdc-1* and *sdc-2* RNAi. (A) Bar chart showing increase in *her-1a* transcript levels as measured by RT-qPCR after treatment with *sdc-2* and *sdc-1* RNAi compared to empty vector (EV) in WT, *xol-1* and *xol-1 sex-1* mutants. *her-1a* upregulation was not statistically significantly different in WT vs *xol-1* (*sdc-2* RNAi p=0.24, *sdc-1* RNAi p=0.59), WT vs *xol-1 sex-1* (*sdc-2* RNAi p=0.07, *sdc-1* RNAi p=0.14), and *xol-1* vs *xol-1 sex-1* (*sdc-2* RNAi p=0.21, *sdc-1* RNAi p=0.33). P-values obtained from paired two-tailed Student’s t-test. Error bars indicate SEM.

**Fig. S4:**
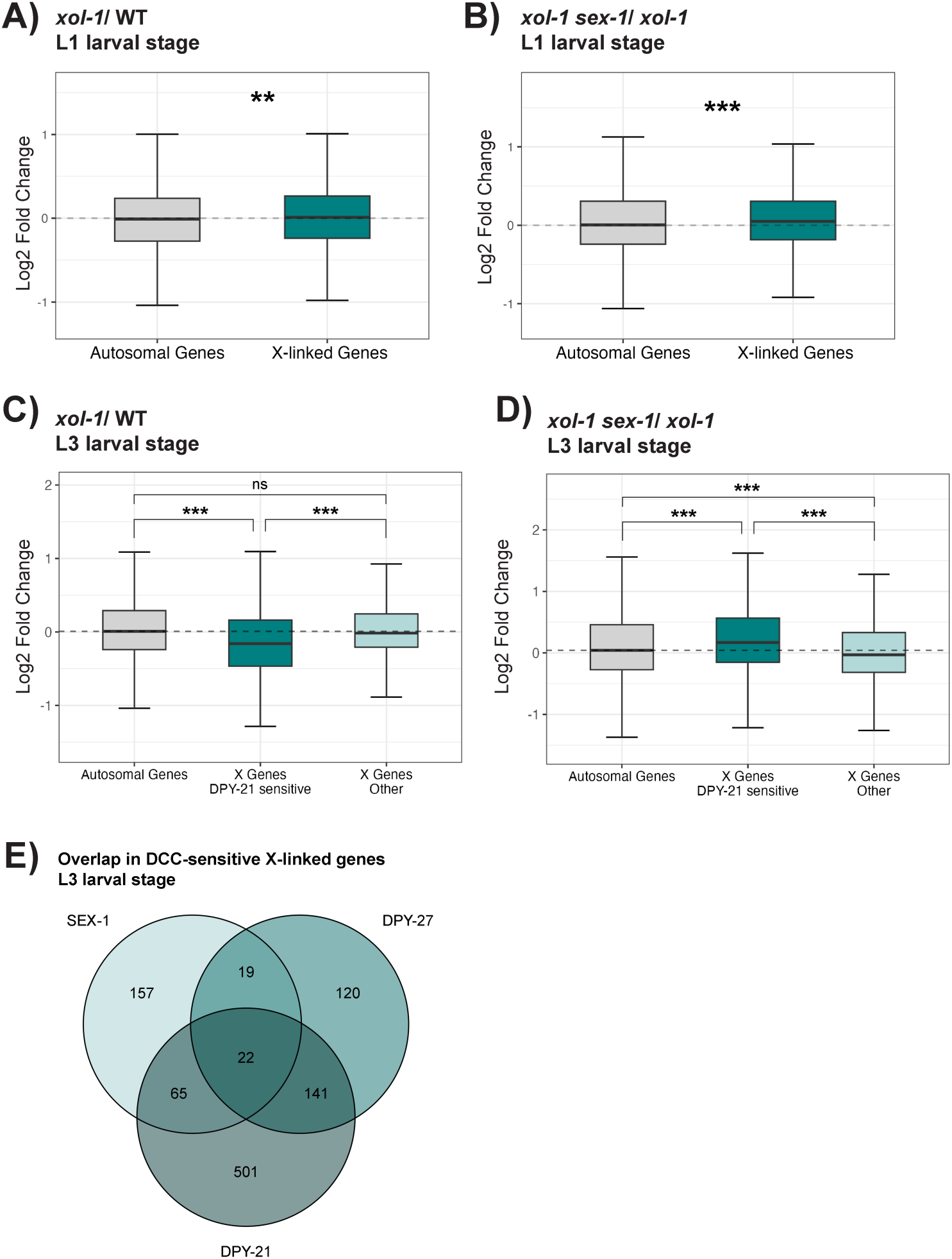
Disrupted X chromosome gene expression in *xol-1* and *xol-1 sex-1*. (A-B) Boxplots showing the distribution of log2 fold change for genes on the autosomes and X chromosomes in (A) *xol-1*/ WT (median log2 fold change (X - A) = 0.02) and (B) *xol-1 sex-1/ xol-1* (median log2 fold change (X - A) = 0.04) at the L1 larval stage. n > 3. Statistical significance was obtained using Wilcoxon rank-sum test. Asterisks indicate level of statistical significance (* p < 0.05, ** p < 0.005, *** p < 0.0005, ns not significant). (C-D) Boxplots showing the distribution of log2 fold change for genes on the autosomes, X-genes sensitive to the loss of DPY-21 function, and X-genes that are not sensitive to the loss of DPY-21 function, in (C) *xol-1*/ WT and (D) *xol-1 sex-1*/ *xol-1*. DPY-21 sensitive X-genes were defined by genes that demonstrate log2 fold change > 1 due to the loss of DPY-21 function. DPY-21 sensitive genes were obtained from analysis of datasets published in Trombley et al. 2024 [43]. Statistical significance was obtained using Wilcoxon rank-sum test. Asterisks indicate level of statistical significance (* p < 0.05, ** p < 0.005, *** p < 0.0005, ns not significant). (E) Overlap in X-linked genes that are upregulated by 2-fold in the absence of *xol-1-*independent SEX-1 function, DPY-27, and DPY-21.

**Table S1:** P-values for experiments quantified in Figure 2, Figure 6, and Figure 7.

**Table S2:** Sequences for primers and CRISPR reagents used in this study

